# Novel sex-biased outcomes in neuroblastoma are associated with distinct gene expression and chromosomal loss patterns

**DOI:** 10.1101/2024.06.14.599094

**Authors:** Vishwa Patel, Mariya Raleigh, Lorraine-Rana E. Benhamou, Xin Shen, Erika C. Espinosa, Niharika R. Badi, Dennis A. Sheeter, John T. Powers

## Abstract

The worst patient outcomes in neuroblastoma are driven by high-risk disease^1,2^, which is divided into similarly sized *MYCN* amplified and *MYCN* non-amplified patient subgroups^3^. Male patients have been reported to have slightly worse outcomes than females in all-patient analyses of multiple studies^3,4^. However, we show here that in *MYCN* non-amplified high-risk and stage 4s low-risk disease, female patients have significantly worse overall survival than males. Female *MYCN* non-amplified high-risk patients highly express *H19* and *DLK1*, both of which drive cell growth *in vitro* and are associated with worse outcomes in females but not males. Further, chromosome-specific expression analysis of these patients reveals broad sex disparities in chromosomal patterning, including female-specific retention of chromosome 11q, a pattern typically reserved for *MYCN*-amplified disease^5,6^. Finally, we show that *H19*, a known *let-7* microRNA target^7^, sequesters *let-7* in females, providing a rationale for worse female survival and reconciling retention of chromosome 11q. We propose that this novel sex-based outcome disparity is driven by *let-7* inhibition, expanding on a model of neuroblastoma development where *let-7* mitigation is central to disease pathology^8^.

## Introduction

Neuroblastoma (NB) is an extracranial solid tumor in children that arises from the neural crest lineage^9,10^. According to the Children’s Oncology Group, NB can be categorized as low-risk (LR), intermediate-risk (IR), or high-risk (HR) based on stage, patient age, favorability of histology, *MYCN* status, and presence of other genetic aberrations such as ploidy or chromosomal loss^11,12^. LR and IR NB have favorable overall survival (90-95%), as do stage 4s tumors, which often regress without intervention^10,13,14^. In contrast, HR disease is notoriously difficult to treat and has an OS rate below 50%^15,16^. Consequently, HR NB patients endure one of the most intensive genotoxic therapeutic regimens in all of oncology^17^. Survivors are often left with life-long health complications and increased risk for secondary cancers. An improved understanding of HR genetic drivers will result in better patient stratification and may lead to the development of more effective, less genotoxic treatments that improve patient survival and post-therapy survivorship. While sex and racial disparities in cancer outcomes have been previously described^3,4,18–21^, frequent all-patient analyses are insufficient for understanding the complex genetic subgroupings of cancers such as NB^3,4,18–21^. Large-scale studies of patient outcomes report that overall, males with NB fair slightly worse than females. However, it is important to consider that the incidence of NB and HR disease are both also higher in males. On a per-patient basis within distinct subgroups, male and female outcomes have not been appropriately investigated, in particular within the HR NB patient population. This concern spans many cancer types as patient studies often lack annotations for categories such as genetic or risk subgroup, sex, or race. A broad approach to comprehensive annotation of cancer patients would facilitate a deeper understanding of the genetic distinctions between subpopulations, enable the identification of more refined subcategories of NB, and lead to improved discovery of novel therapeutic targets.

HR NB is equally subdivided into either *MYCN*-amplified(*M-Amp)* or *MYCN* non-amplified (*M-Non)* HR subgroups. *MYCN* status is also associated with common chromosomal aberrations, such as deletions of 1p, 11q, 14q, and 17q gain^5,6,22,23^. Further, *MYCN* mRNA is targeted by the *let-7* tumor suppressor microRNA family^24^. We previously showed that high levels of *MYCN* mRNA in *M-Amp* disease can turn the tables on *let-7*, where the 3’ UTR of *MYCN* mRNA efficiently sequesters *let-7* and inhibits its function^8,25^. This discovery also explained the longstanding observation that chromosome (Chr) 11q, known to harbor a highly expressed *let-7* locus (*let-7a-2*), is frequently lost in *M-Non* but not in *M-Amp* disease. This work established that increased levels of Chr 11q-derived *let-7a* are functionally impaired by *MYCN* mRNA. Further, this was the first demonstration that noncoding elements within a protein-coding mRNA can independently contribute to both oncogenicity and selective genetic patterning in cancer.

Here, we assessed potential sex and race biases in NB. We discovered a striking female-directed bias in *M-Non*/HR NB patients that associates with high expression of *H19* and *DLK1* ^26–29^. Biased outcomes were also identified in *M-Non* stage 4s disease, as well as an opposing bias affecting male African American patients. We identified significant differences in chromosomal loss patterns between female and male *M-Non*/HR NB patients using a novel analytic approach to differential gene expression. Finally, we connected these sex-biased outcomes and chromosomal loss patterns with *let-7* sequestration by *H19*, expanding on our previous report on the intersection between *MYCN* expression, *let-7* inhibition, and chromosomal patterning in *M-Amp* disease^8,25^.

## Results

### Female neuroblastoma patients have worse overall survival in High-Risk MYCN Non-Amplified disease

Overall, NB incidence and the occurrence of HR disease are higher in males than in females^3,4,30^. However, an in-depth analysis of female *vs.* male outcomes in HR disease has yet to be reported. We first assessed potential sex bias in multiple NB clinical subsets through analysis of a 498-patient cohort (SEQC) with sufficient annotation to simultaneously interrogate risk stratification, *MYCN* status, and patient sex^31^. No difference in overall survival (OS) between the sexes was observed in this and two additional datasets (TARGET, Kocak) when risk status was not considered (***Extended Data Table 1, Extended Data Fig. 1a, 1b, 1c***)^32,33^. SEQC patients with LR disease also exhibited no significant difference in survival between males and females (***Figure 1a***, top right panel). However, among HR patients, females displayed a striking pattern of significantly worse OS (*p = 0.0075*, ***Figure 1a***, top left). Analysis of the *M-Amp* and *M-Non* subgroups within HR disease revealed significantly worse OS in female *M-Non*/HR patients (*p = 0.021*), as opposed to no difference in outcomes in the *M-Amp*/HR subgroup (***Figure 1a***, lower right and left panels, respectively). We next sought to validate this finding within additional datasets. The Kocak NB patient study has annotations for *MYCN* status, patient age, stage, and sex but not risk status (***Extended Data Table 1***). Therefore, we considered stage 3 or stage 4 patients older than 18 months (approximation of HR disease)^4^, again revealing that higher-risk, late-stage female NB patients had significantly worse OS (*p = 0.0037,* **Extended *Data*** Fig. 2a). We then assessed female and male OS in the TARGET NB study (***Extended Data Table 1***). An initial comparison of HR patients yielded no difference in outcomes between female vs. male patients (***Extended Data*** Figure 2b). Whereas the Germany-based Kocak study has a strong Caucasian patient bias (*personal communication*), African American (AA) patients represent the second largest group in the TARGET study (over 12% of subjects). We reasoned that the increased diversity within the TARGET study may have contributed to our initial failed validation of female sex bias. Upon exclusion of AA patients from our analysis, we again observed the same striking pattern of worse OS in female *M-Non*/HR patients as in the SEQC study (***Extended Data*** Fig. 2c). When analyzed in isolation, the AA patient group (n = 31) exhibited an inverse pattern, where male HR patients experienced dramatically worse OS than females (***Figure 1b***).

**Figure 1.**
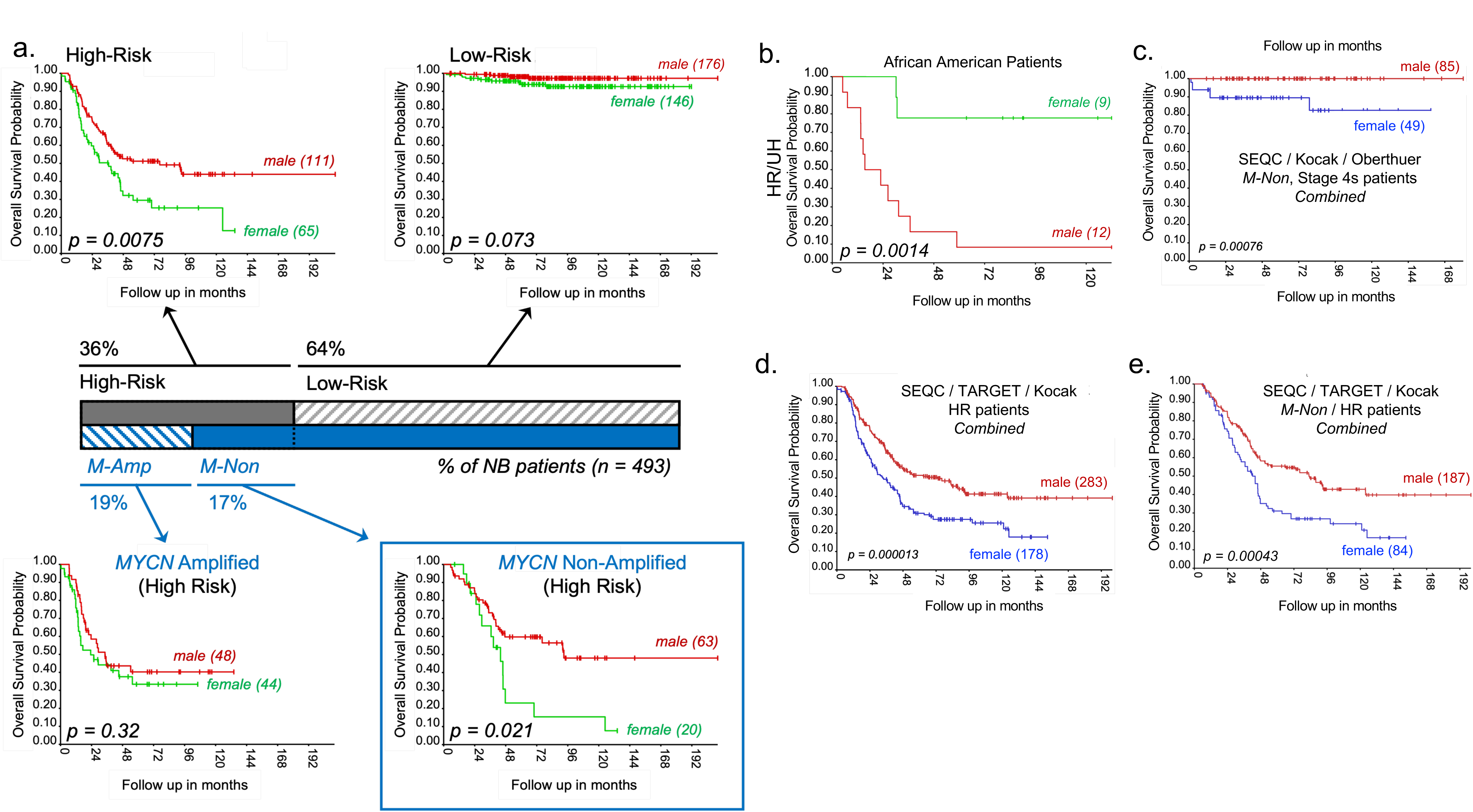
Identification of broad sex-biased outcomes in neuroblastoma. **a)** Kaplan-Meier (KM) analysis of overall survival (OS) in male vs. female neuroblastoma (NB) patients relative to *MYCN* and risk status. Center bar represents NB patients from the SEQC 498 dataset; HR (n=176), LR (n=322), *M-Amp*/HR (n=92), and *M-Non*/HR (n=83). Data source: SEQC 498 (GSE62564). **b)** KM analysis of OS in female and. male African American high- risk//unfavorable histology NB patients. Data source: TARGET 249 (R2, Asgharzadeh). **c)** KM analysis of OS in male vs. female *M-Non*/stage 4s NB patients. Combined data source: SEQC 498 (GSE62564), Kocak 649 (GSE45547), Oberthuer 251 (E-TABM-38). **d)** KM analysis of OS of male vs. female HR NB patients. Combined data source: SEQC 498 (GSE62564), Kocak 649 (GSE45547; approximation of high-risk constituting stage 3 and 4 patients >18 months old), TARGET 249 (R2, Asgharzadeh). **e)** KM analysis of OS of male vs. female *M-Non*/HR NB patients. Combined data source: SEQC 498 (GSE62564), TARGET 249 (R2, Asgharzadeh), Kocak 649 (GSE45547). All *p* values were determined by log-rank test.

This striking yet opposite-sex bias in OS led us to evaluate other subgroups within NB. In particular, analysis of *M-Non* stage 4s patients, whose prognosis is overall favorable, also revealed a significant difference in OS between females and males. When analyzed individually, females in the SEQC and Kocak cohorts had significantly worse OS than male *M-Non* stage 4s patients (*p = 0.019, 0.021*, respectively**, *Extended Data*** Fig. 3a***, 3b***). A third study revealed a similar trend but did not achieve significance (*p = 0.*17, **Extended *Data*** Fig. 3c). Combined analysis of these three studies revealed a strong sex bias, where female OS was much worse than male 4s patients (*p = 0.*00076, **Figure *1c***). Indeed, the only patients who died from disease within this group were female (8/49), whereas there were no male deaths (0/85) (***Figure 1c, Supporting Data file 1***). This is surprising, given that stage 4s disease has an excellent prognosis, with over 90% survival and close to 50% spontaneous regression^34^. Our results further suggest a possible 100% survival rate in *M-Non* stage 4s male patients, compared to 16.4% death in females (***Figure 1c***). We next combined higher risk patients from the SEQC, Kocak, and TARGET (AA patients excluded) studies to increase sample size and power for the previously identified sex bias in *M-Non*/HR disease. From this analysis, we observed an increased statistical significance associated with adverse female-biased OS in HR disease (n = 461 patients, *p = 1.3 x 10^-^*^5^, ***Figure 1d, Supporting Data file 1***) and a stronger representation of sex-biased OS in female *M-Non*/HR disease (n = 271, *p = 4.3 x 10^-4^,* ***Figure 1e, Supporting Data file 1***).

### H19 and DLK1 are associated with poor survival in female, but not male, neuroblastoma patients

Differential gene expression analysis of the top 500 genes in female (n = 14) *vs.* male (n = 26) *M-Non*/HR dead from disease patients (SEQC) revealed an expected strong shift towards female enrichment in x-y plots of gene expression (***Figure 2a***, right panel). In contrast, this geneset did not exhibit enrichment among female patients in *M-Amp*/HR dead from disease nor *M-Non*/LR living clinical subgroups, suggesting that female *M-Non*/HR patients have a distinct gene expression signature (***Figure 2a***, middle and left panels). Among the top twenty genes upregulated in female *M-Non*/HR patients, the long noncoding RNA *H19* and delta-like non-canonical Notch ligand 1 (*DLK1*) emerged as the most upregulated non-Chr X genes (***Figure 2a***, right panel, ***Extended Data*** Fig. 4a left panels***, Supporting Data file 2***). Further, *H19* and *DLK1* expression is significantly correlated in females (r = 0.52, p = 0.019) but not male *M-Non*/HR patients, suggesting possible co-regulation in female patients (***Figure 2c, Supporting Data file 2***). A similar analysis of HR- approximated patients in the Kocak study again showed that *H19* and *DLK1* are significantly upregulated in female patients (***Figure 2b, Extended Data*** Fig. 4a right panels***, Supporting Data file 2***). Further, despite very small patient numbers (n = 5 female, 6 male), a comparison of *H19* and *DLK1* expression in *M-Non* stage 4s patients with progressive disease shows higher, although not significant, expression in female patients (***Extended Data*** Fig. 4b). Of note, expression of *H19* and *DLK1* in the adrenal medulla is among the highest for all tissues (***Extended Data*** Fig. 5), introducing the possibility that high *H19* and *DLK1* expression in female *M-Non*/HR patients may represent retained expression from a common NB tissue of origin. Cox-regression analysis of OS Hazard Ratios of the top female-enriched non-X Chr genes reveals that *DLK1* and *H19* (and its embedded microRNA, *miR-675*) are significantly associated with worse outcomes in only female patients (***Figure 2d, Supporting Data file 2***). This association was validated through Kaplan-Meier analysis of *M-Non* patients in the SEQC and Kocak studies (***Extended Data*** Fig. 6a***, 6b***). No association with female or male outcomes was noted in *M-Amp* disease patients from either study (***Extended Data*** Fig. 7).

**Figure 2.**
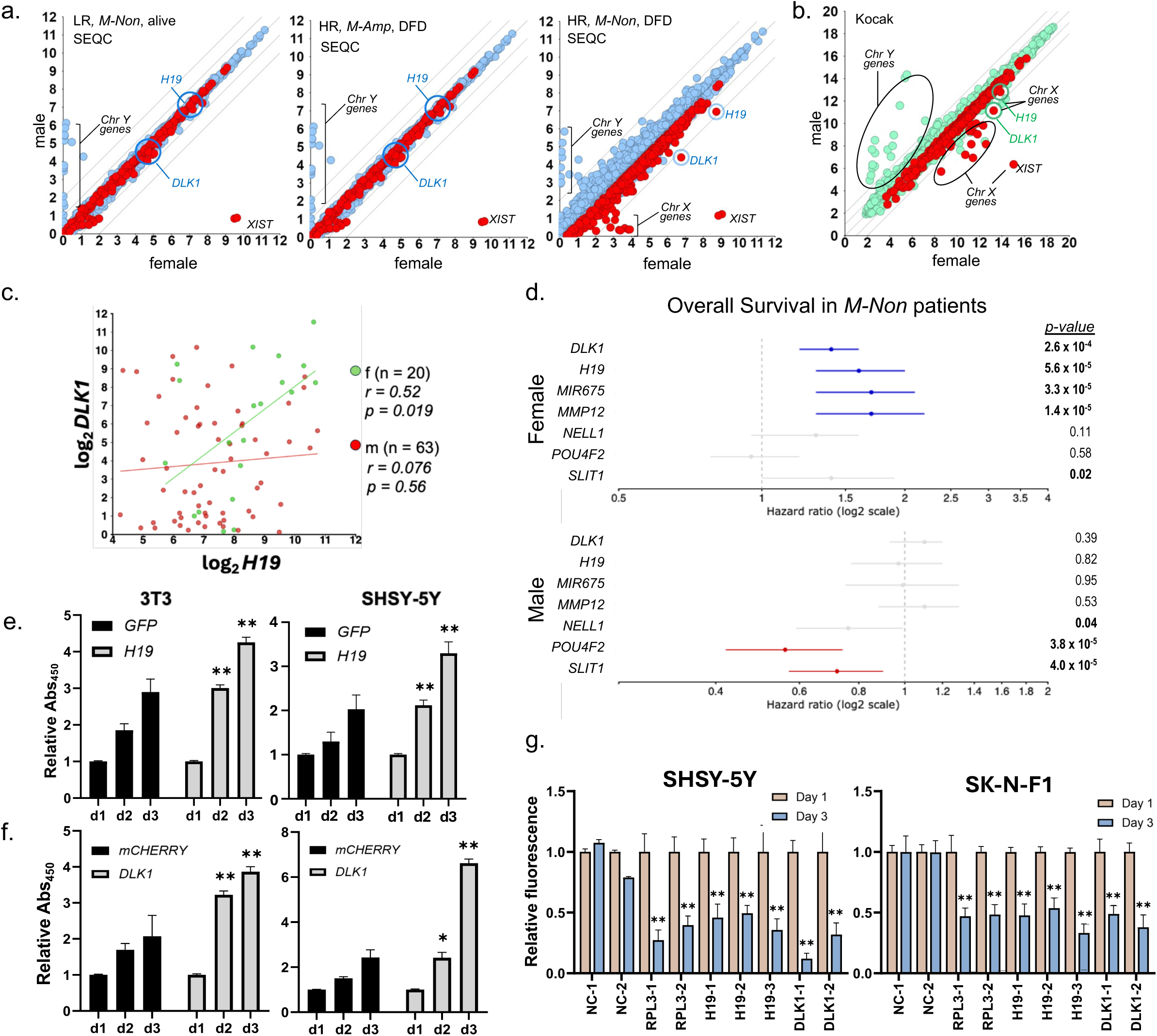
*H19* and *DLK1* expression differences in male vs. female neuroblastoma patients. **a)** X-Y plot of differential expression analysis of male vs. female NB patients. Red dots = top 500 genes most significantly upregulated in LR/*M-Non/*alive patients (left); HR/*M- Amp/*DFD patients (center); HR/*M-Non/*DFD patients (right). Data source: SEQC 498 (GSE62564). **b)** X-Y plot of differential expression analysis of male vs. female st4/*M-Non* NB patients. Red dots = top 500 significantly upregulated genes in (2a). Data source: Kocak 649 (GSE45547). **c)** X-Y plot of gene expression correlation between *H19* and *DLK1* expression levels in male and female HR/*M-Non*/DFD patients. Green line, f = females, red line, m = males. p-values determined by ANOVA. Data source: SEQC 498 (GSE62564). **d)** Cox proportional hazard regression analyses for female vs. male *M-Non* NB patients (as labeled). Selected genes are most significantly upregulated non-X Chr-associated genes in *M-Non* NB patients. Extended lines from central dots indicate 95% CI. Blue line = significant correlation > 1.0, red line = significant correlation < 1.0. Data source: SEQC 498 (GSE62564). **(e)** and **(f)** Colorimetric growth assays on 3T3 and SHSY-5Y cells relative to GFP construct-based expression of *H19* and *DLK1*. Significance determined by two-way ANOVA, followed by Dunnett’s multiple comparison test; bars represent average +/- SEM; *p<0.05, **p<0.01, <0.001, <0.0001. **g)** CRISPR/Cas9 mediated gene knockout of *H19* and *DLK1* in SHSY-5Y and SK-N-F1 cells. Cells were transfected on d0 with control, *DLK1* and *H19* gRNAs. Loss of *H19* and *DLK1* targeted with gRNAs against negative control gRNA were measured at d1 and d3 via flow cytometry. Significance determined by two-way ANOVA, followed by Sidak’s multiple comparison tests; bars represent mean+/-SEM; *p<0.05, **p<0.01, <0.001, <0.0001.

*H19* and *DLK1* have been implicated in multiple cancers^26–29^. To assess their ability to promote NB tumor growth, we over-expressed both in murine 3T3 and human *M-Non* SH-SY5Y NB cells. Expressing *H19* or *DLK1* resulted in higher growth rates compared to GFP control cells (***Figure 2e, 2f***). Use of *H19*- or *DLK1*-targeting small interfering RNAs (siRNAs) individually or in tandem in SH-SY5Y cells resulted in significant growth inhibition (***Extended Data*** Fig. 8a***, Supporting Data file 3)***. We next used a Cas9-mediated gene disruption growth assay modeled on classical gRNA dropout screens to disrupt *H19* and *DLK1*. Both SH-SY5Y and SK-N-F1 cells displayed significant reductions in fluorescence driven by positive control *RPL3* and both *H19* and *DLK1* gRNAs (***Figure 2g, Extended Data*** Fig. 8b ***Supporting Data file 2***). These data suggest that *H19* and *DLK1* contribute to aggressive tumor growth, thereby contributing to the poor outcomes we report in *M-Non*/HR female patients.

### Differential gene expression analysis reveals striking sex-biased chromosomal loss patterns

When we assessed the genomic location of the top 500 genes significantly increased in female *M-Non*/HR patients, we identified a strong enrichment for genes residing in the q arm of Chr 11 (15.8%). Female-biased gene expression was also noted for Chr 4 (10.8%) and Chr 12q (10.2%) (***Figure 3a***, *bottom panel, **Supporting Data file 4***). This prompted us to examine the chromosomal distribution of male-enriched genes, where we observed an overrepresentation of Chr 1p (10.6%) and Chr 19 (11.4%). (***Figure 3a*,** *top panel, **Supporting Data file 4***). We reasoned that these enrichment patterns may reflect chromosomal patterning differences between female and male *M-Non*/HR patients. To examine this directly, we first assessed differential expression patterns in *M-Amp* vs. *M-Non* NB patients, where chromosomal patterns are well established. First, looking at Chr 13, not known to have a Chr loss signature in NB, we observed no significant difference in gene expression by volcano plot and paired t-test (***Figure 3b***). Chr 1p, however, known to be lost in almost all *M-Amp* disease ^6,35^, displayed a significant shift of gene expression toward the *M-Non* group (*p = 8.4 x 10^-^*^100^, ***Figure 3c, Supporting Data file 4***). Similarly, Chr 11q, known to be frequently lost in *M-Non* disease^5,6,8,22^, exhibited an opposite shift in gene expression towards *M-Amp* disease (*p = 0.0028,* ***Figure 3d, Supporting Data file 4***). Our analytical approach successfully recapitulated known chromosomal patterning between patient subgroups, suggesting that this method can be used to identify novel chromosomal abnormalities.

**Figure 3.**
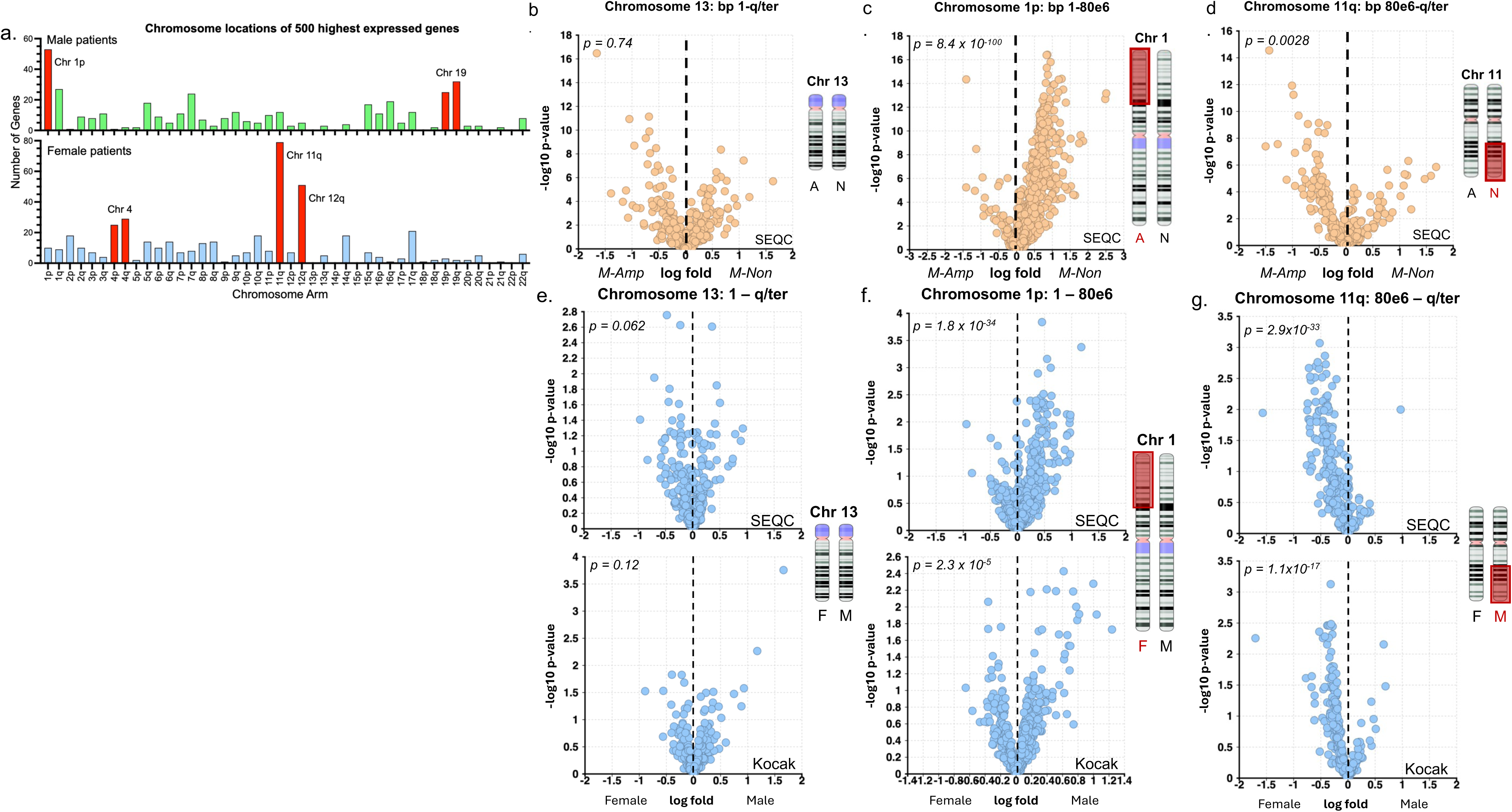
Chromosome-associated differences in gene expression in male vs. female neuroblastoma patients. a) Chromosomal location of the 500 most highly expressed genes in female vs. male HR/DFD NB patients. Red bars: >20 genes present on respective chromosomes. Data source: SEQC 498 (GSE62564, n=40, composed of 14 females and 26 males). **b), c), d)** Volcano plot of Chr 13, Chr 1p, and Chr 11q genes expression in *M-Amp*/HR and *M-Non*/HR NB patients (n=91). Data source: SEQC 498 (GSE62564) **e), f),** and **g)** Volcano plot of Chr 13, Chr 1p, and Chr 11q genes expression in female and male NB patients. Data source: SEQC 498 (top plots; GSE62564; *M-Non*/HR/DFD subset; n=40) and Kocak 649 (bottom plots; GSE45547; st4 patients >18 months old subset; n=94).

Chr 13 genes were not enriched in either female or male patients (1% and 0.6%, respectively (***Figure 3a***, ***Supporting Data file 4)***. Accordingly, we observed no significant sex bias in Chr 13 gene expression in patients from either the SEQC or Kocak studies (***Figure 3e***). Next, we examined Chr 1p and observed a significant shift toward male patients in both the SEQC and Kocak studies (*p = 1.8 x 10^-^*^34^*, 2.3 x10^-^*^5^*, respectively,* ***Figure 3f, Supporting Data file 4***). Further, analysis of Chr 11q genes showed a significant shift in gene expression towards female patients in both studies (*p = 2.9 x 10^-^*^33^*, 1.1 x 10^-17^, respectively,* ***Figure 3g, Supporting Data file 4***). The other chromosomal enrichment patterns identified across Chr 4, Chr 12q, and Chr 19 (***Figure 3a***) also showed significant sex-biased shifts (***Extended Data*** Figure 9***, Supporting Data file 5***). These results demonstrate for the first time that sex-biased patterns of chromosomal loss exist in NB. While this analysis does not allow for *de novo* determination of directionality for a given chromosomal imbalance, Chr 1p and 11q aberrations are well-known in NB. The patterns we observed are therefore likely due to chromosomal arm loss within the deficient group^5,6,22,36–38^. Chr 4^36,37,39^ and Chr 12q^40–42^ losses have been associated with poor outcomes in NB, which is consistent with the female-biased poor OS we observed in *M-Non*/HR disease. Whole Chr 19 gain^36^ has been associated with better outcomes, suggesting that we may have identified a pattern of Chr 19 gain in males as opposed to loss in females.

Finally, we examined the local genomic regions near both *DLK1* and *H19.* We observed that the Chr 14q region near the *DLK1* locus is associated with a female-enriched shift, suggesting a genetic dose difference favoring females for that region (***Extended Data*** Figure 10a***, Supporting Data file 6)***. *H19*, however, showed no significant shift in either direction (***Extended Data*** Figure 10b ***Supporting Data file 6***). This discrepancy suggests that *DLK1* elevation in females may be due, at least in part, to differences in gene copy number. In contrast, elevated *H19* expression may be through a more direct mechanism in females.

### H19 inhibits let-7 in female M-Non/HR patients in a manner similar to MYCN mRNA in M-Amp disease

Despite the difference in *MYCN* amplification status, female *M-Non*/HR patients have similar Chr 1p and Chr 11q patterns to *M-Amp* patients (***Figure 3f, 3g***). To explore this further, we repeated our Chr-specific differential gene expression analysis of Chr 13, Chr 1p, and Chr 11q in male and female NB patients. *M-Amp* male patients displayed the expected pattern, revealing loss of Chr 1p and retention of Chr 11q compared to the *M-Non* group (***Figure 4a***, upper panels, ***Supporting Data file 7***). Female patients displayed the expected patterns for Chr 13 and Chr 1p, but diverged for Chr 11q, showing a mildly significant shift towards *M-Non* patients. This suggests that female *M-Non* patients may indeed retain Chr 11q more than male patients, further supporting the idea that *M-Non*/HR females may display a classically *M-Amp* loss pattern for Chr 1p and Chr 11q.

**Figure 4.**
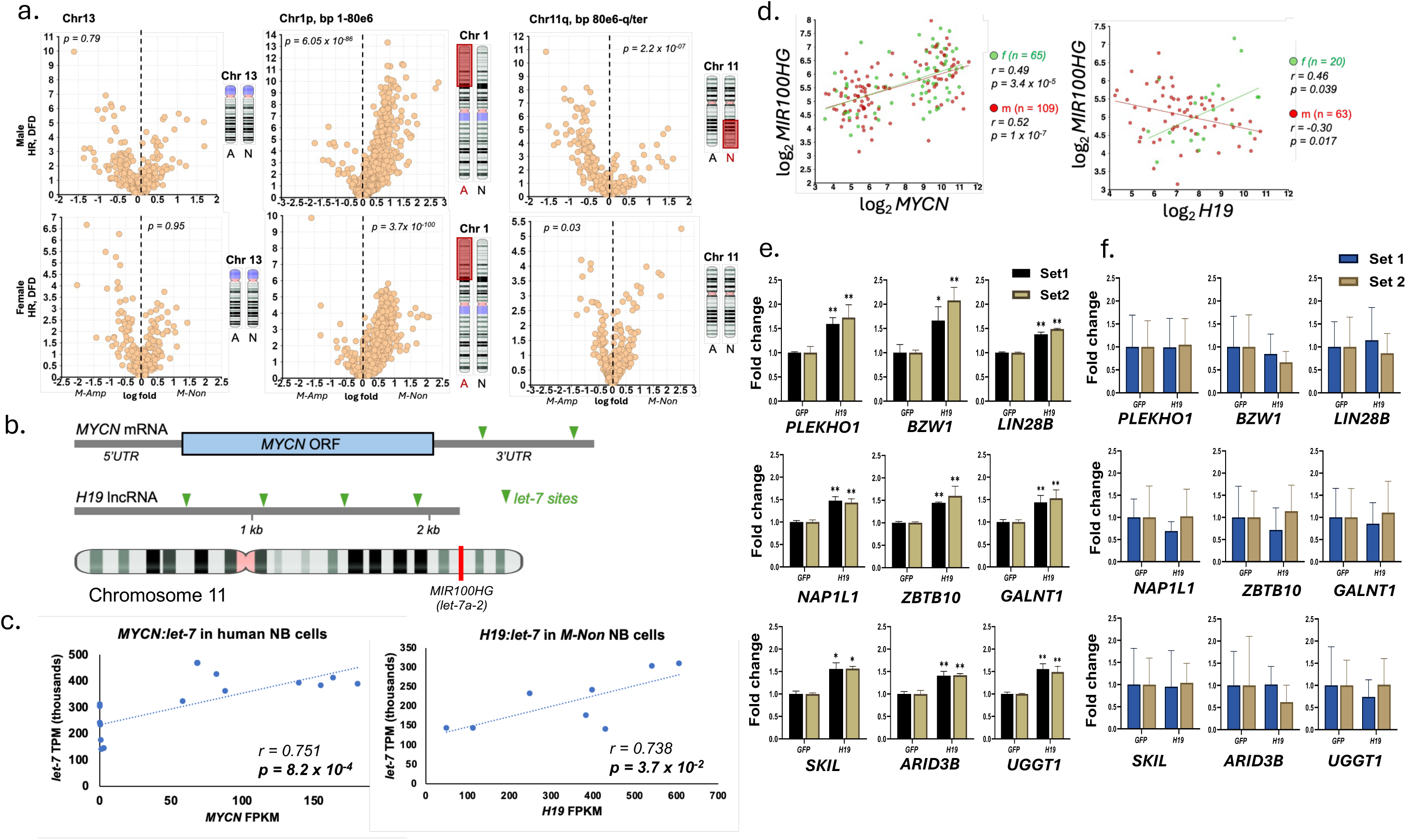
*H19* is strongly associated with let-7 expression in neuroblastoma. **a)** Volcano plots of chromosomal-linked gene expression HR dead from disease NB patients stratified by *MYCN* status and sex. Top row: plots of male *M-Non* (n=26) and *M-Amp* patients (n=26) for stated chromosomal regions; Bottom row: plots for female *M-Non* (n=14) vs. *M-Amp* (n=25). Yellow dots: genes stemming from specified chromosomes in male or female; red box over chromosome = identified relative chromosome deficiency. Data source: Data source: SEQC 498 (GSE62564). **b)** Schematic of *MYCN* mRNA and *H19* lncRNA *let-7* sites. Green triangles = *let-7* miRNA target sites. Bottom: Location of let- 7a-2 host gene (MIR100HG) on chromosome 11. **c)** Correlation plots of *MYCN* vs. *let-7* expression in human *M-Non* and *M-Amp* NB cell lines (n=16; left) and *H19* vs. *let-7* expression in human *M-Non* NB cell lines (n=8; right) were derived from NGS. Data sets will be available under the GEO reference series GSE (http://www.ncbi.nlm.nih.gov/geo/query/acc.cgi?acc=GSE (TBD1, TBD2)). P-values were derived by Pearson Product Moment Correlation (PPMC) from correlation coefficient (*r*). **d)** X-Y plots of gene expression correlation between *MYCN* and *MIR100HG* (left) in HR NB patients (left), *H19* and *MIR100HG* in *M-Non*/HR NB patients (right). Green line, f =females; red line, m =males. Data source: SEQC 498 (GSE62564; n=176); *p<0.05, **p<0.01; p-values determined by ANOVA. ***e*)** Expression levels of *let-7* target genes in *M-Non* SHSY-5Y cells expressing *H19. **f)*** Expression levels of *let-7* target genes in *M-Amp Be(2)c* cells expressing H19. e),f) Significance determined by two-way ANOVA, followed by Sidak’s multiple comparison tests; bars represent average +/- SEM; *p<0.05, **p<0.01, ***<0.001, ****<0.0001.

We previously established that in *M-Amp* NB, *MYCN* mRNA, which is a *let-7* microRNA target (***Figure 4b***), sequesters *let-7* in *M-Amp* disease. This functional mitigation relieves the selective pressure to lose Chr 11q, itself housing one of the highest expressed *let-7* loci, *let-7a-2*^8^ (***Figure 4b, Extended Data*** Figure 11a***, 11b***). This work provided a mechanism by which Chr 11q is retained in *M-Amp* NB but preferentially lost in *M-Non* disease and also established that noncoding elements within a protein-coding mRNA can have autonomous oncogenic function with genetic consequences. Our current observation, where female *M-Non*/HR patients have a Chr 11q pattern that matches classical M-Amp patterning, may have consistencies with this original discovery. *H19*, which is highly expressed in these female patients (***Figure 2a, Extended Data*** Figure 4), is known to contain four *let-7* sites (***Figure 4d***)^7^ and has been reported to sequester *let-7*^43–46^. *H19* shares several characteristics in NB with *MYCN*, a known *let-7* sponge. As expected, *MYCN* correlates with *let-7* levels across *M-Amp* and *M-Non* NB cell lines (***Figure 4c****, left panel, **Supporting Data file 7***)^8^. Similarly, *H19* correlates with *let-7* levels in *M-Non* NB cell lines (***Figure 4c****, right panel*). This relationship holds relative to the *let-7a-2* host gene, *MIR100HG*, in human NB patients, where *MYCN* levels correlate with *MIR100HG* in HR NB patients (***Figure 4b*** and ***4d****, left panel*). *H19* levels also positively correlate with *MIR100HG* expression in *M-Non* HR disease, but only in females (***Figure 4d****, right panel*). This suggests that *H19* may play a role in *let-7* inhibition in only female *M-Non*/HR patients, uniting the observed sex-based disparity in OS with Chr 11q (*let-7a-2*) patterning in *M-Amp* and female *M-Non* patients.

To test this more directly, we examined the expression levels of nine sensitive *let-7* targets in *H19* overexpressing SH- SY5Y cells. Compared to GFP, *H19*-expressing cells showed significantly elevated expression of all *let-7* targets (***Figure 4e, Extended Data*** Figure 11d***, Supporting Data file 7***), which is consistent with *H19* sequestration of *let-7*. When we repeated this analysis in *M-Amp* BE(2)C cells, where established *let-7* sponge *MYCN* is expressed at very high levels ^8^, we failed to observe significant enrichment of any *let-7* target (***Figure 4f, Supporting Data file 8***). This result is not unexpected, given that *let-7* is already sequestered and functionally suppressed by *MYCN* mRNA in these cells, making expression of *let-7*-interacting *H19* redundant and minimally impactful in these cells. Consistent with these results, female *M-Non*/HR NB patients have significantly enriched expression of *let-7* target mRNAs containing two or more *let- 7* sites (*n = 133 let-7 targets, **Extended Data*** Figure 11c***, Supporting Data file 9***). We next examined the effects of *H19* depletion on *let-7* target mRNAs. Upon siRNA-mediated knockdown of *H19* in *M-Non* CHLA-42 NB cells (***Extended Data*** Figure 12a), we observed significantly reduced expression of most *let-7* target mRNAs (***Extended Data*** Figure 12b***, Supporting Data File 10***). These results suggest that *H19* sequesters and inhibits *let-7* in female *M-Non*/HR patients in a manner similar to *MYCN* mRNA in *M-Amp* disease^8^ and further provide a plausible rationale for worse female outcomes. This novel role for *H19* in female NB patients provides a selective mechanism by which these female patients tend to retain Chr 11q, a pattern previously only associated with *M-Amp* patients.

## Discussion

Prior to this work, male and female HR NB outcomes had not been directly investigated. Understanding the origins and driving mechanisms behind the sex-based disparity in OS described here may lead to an improved understanding of female-specific cancer genetics and sex-specific approaches to therapy, a first for NB. While primarily male-biased, worse outcomes have been reported in over twenty cancer types^47,48^, the majority of these disparities were identified by looking at all patients within a given cancer^48^. Given the results of our compartmentalized analysis, we predict that sex biases are even more widespread in human cancer, that these biases will be more divergent in some patient subgroups, and that the directionality may change depending on the subgroup in question. The results of our analysis provide a rationale for widespread sub-group analysis across most if not all, tumor types. Therefore, it is imperative to consider tractable subgroups when analyzing patient outcomes. To facilitate such subgroup-focused analysis, we must strive to maximize annotation when performing large-scale patient studies. Of eight patient datasets available for possible outcomes analysis on the r2 database, only four listed patient sex within their available annotations^49^. In contrast, the TARGET study with only 249 patients was annotated to the degree that it included patient race, allowing us to discover an additional and opposite-sex bias in African American patients (***Figure 4***).

Both *H19* and *DLK1* have been implicated as oncogenes in multiple cancers^26–29,50–54^. *H19* is a long non-coding RNA with known functions that include regulation of gene expression and microRNA sequestration^50,55^. *DLK1*, a transmembrane protein involved in the highly conserved NOTCH signaling pathway, can exert control over cell-to-cell communication, proliferation, and differentiation^26,27^. We found similar and robust pro-growth functions for both *H19* and *DLK1* in murine 3T3 embryonic fibroblasts and in human *M-Non* SH-SY5Y NB cells (***Figure 2e***, ***2f***, ***2g***, and ***Extended Data*** Figure 8a). The local organization of the *H19* (*h*) and *DLK1* (*d*) loci are similar. They both reside in imprinted regions, and both loci are comprised of a protein-coding gene (*DLK1*(*d*) or *IGF2*(*h*), both imprinted), followed by long noncoding RNA (*MEG3*(*d*) or *H19*(*h*), both imprinted), followed by another protein-coding gene (*RTL1*(*d)* or *MRPL23*(*h*))^56,57^. These similarities suggest the possibility of shared regulation.

NB is a disease of genetic compartmentalization, both in terms of expression patterns of key genes and repeating patterns of chromosomal gains and losses. We identified a surprising sex disparity in chromosomal imbalance across multiple chromosomes, including Chr 1p and Chr 11q (***Figure 3a***, ***3f***, and ***3g***). This observation revealed an unexpected alignment between female *M-Non*/HR and classical *M-Amp* patterning as both groups appear to lose Chr 1p and retain Chr 11q. Our previous work established that in *M-Amp* NB, *MYCN* mRNA, which contains two *let-7* sites, efficiently sequesters *let-7* and inhibits its function^8,25^. This work provided the first plausible explanation that Chr 11q, which harbors one of the most highly expressed *let-7* loci (*let-7a-2*), is frequently lost in *M-Non*, but not in *M-Amp* disease. We previously demonstrated that noncoding elements with an oncogenic mRNA can independently contribute to both tumor pathology and selective genetic patterning in cancer^8^. The alignment between female *M-Non*/HR and *M-Amp* patterns led us to discover that *H19*, known to have four *let-7* sites^7^, sequesters *let-7* in female *M-Non*/HR patients in a manner mirroring that of *MYCN* in *M-Amp* disease (***Figure 4, Extended Data*** Figure 11c***, 12***). This observation provides a plausible rationale for the female retention of Chr 11q in *M-Non*/HR disease. The mechanistic similarity of noncoding *H19* and *MYCN* mRNA function validates our earlier work. It suggests that sequestration of *let-7* by either 3’UTRs of protein-coding mRNAs or *let-7* sites within long noncoding RNAs may be a fundamental mechanism of *let-7* mitigation in NB and other cancers.

We show here significant sex- and race-based disparities in OS relative to *MYCN* status in NB through analysis of multiple publicly available patient datasets with good to excellent patient annotations. This discovery provides a novel way to consider how incidence and genetic patterning intersect with disease pathology and patient outcomes in a genetically and clinically heterogeneous disease like NB. Our findings demonstrate the value of highly compartmentalized cohort analysis of large, well-annotated patient datasets. Further, given the frequency and consistency of multiple genetic events in NB, these findings establish a novel paradigm of sex-biased outcomes and chromosomal patterning in NB and solidify the centrality of *let-7* mitigation, which has broad implications for human malignancy.

## Methods

### R2 database

Human NB patient RNA-seq, microarray, and overall survival data sets were obtained from the r2: microarray analysis and visualization platform (http:// r2.amc.nl)^49^. The following data sets were used for analysis: SEQC498 (GEO accession number GSE62564)^31^, Kocak649 (GEO accession number GSE45547)^32^, OBERTHUER251 (E- TAM-38), TARGET249 (r2 internal identifier: ps_avgpres_targetnrbl249_huex10t); results from analysis of the TARGET249 dataset are based upon data generated by the Therapeutically Applicable Research to Generate Effective Treatments (https://www.cancer.gov/ccg/research/genome-sequencing/target) initiative (phs000467)^33^.

### Secondary analysis

Gene expression graphs, volcano plots, Cox regression analysis, and Kaplan–Meier curves analysis of human NB patients were generated using the data analysis tool suite on the r2: Genomics Analysis and Visualization Platform. Data sets used were listed in the R2 database section of this document. Higher-risk NB patients from the Kocak 649 (GSE45547) study were identified as patients with stage 4 disease, >18 months at diagnosis, and patients of any age and stage with *M-Non* tumors. Shifts in differentially expressed chromosome-specific genes were represented as volcano plots for each chromosome. Volcano plots of chromosomal gene expression changes were generated using gene lists from the UC Santa Cruz genome browser. Expression patterns of these gene lists were then obtained from both SEQC-498 (GSE62564) and Kocak 649 (GSE45547) studies on the r2 platform for NB subgroup comparisons.

### Cell Culture

The neuroblastoma cell lines, including CHLA-15 (RRID CVCL_6594), CHLA-20 (RRID CVCL_6602), KAN (SMS-KAN, RRID CVCL_7131), KANR (SMS-KANR, RRID CVCL_7132), KCN (SMS-KCN, RRID CVCL_7133), KCNR (SMS-KCNR, RRID CVCL_7134), SKNF1 (SK-NF-1, RRID CVCL-1702), and BE(2)C (SK-N-BE(2)C, RRID: CVCL_0529) were obtained from the Childhood Cancer Repository at the Texas Tech University Health Science Center, a partner of the Children’s Oncology Group. Additionally, BJ fibroblasts (NHF, ATCC CRL-2522), SH-SY5Y (ATCC CRL-2266), and3T3 (M-MSV-BALB/3T3ATCC CCL-163.2) were acquired from ATCC. All cell lines were cultured in RPMI-1640 media supplemented with 10% heat-inactivated fetal bovine serum (HI-FBS). Cells were incubated at 37°C under 5% CO_2_ and routinely tested negative for mycoplasma contamination. All cell lines utilized are not considered commonly misidentified.

### RNA-Seq

Total RNA was submitted to LC Sciences (Houston, TX, USA) for library prep and sequencing. Sequencing (without spike-ins) was performed at a depth of 40 million 150 bp paired-end reads. Reads were normalized as Fragments Per Kilobase of transcript per Million mapped reads (FPKM). RNA-Seq data sets are (will be) available under the GEO reference series GSE). CHLA-15, CHLA-20, KAN, KANR, KCN, KCNR, CHLA-42, and SKNF1 were submitted for sequencing in biological duplicate.

### Plasmids and Stable Transfection Protocol

Control *GFP* (pPB[Exp]-Puro-EF1A>EGFP); *H19* pPB [ncRNA]-EGFP- EF1A>hH19[NR_002196.2]; and *DLK1* (pPB[Exp]-mCherry-SFFV>hDLK1[NM_003836.7]) vectors were designed and purchased from vectorbuilder.com. 3T3 and SH-SY5Y cells were co-transfected with both 1 µg of desired plasmid and 0.5 µg of hyPBase transposase-expressing vector (pRP[Exp]-mCherry-CAG>hyPBase), following the Lipofectamine™ 3000 (Invitrogen^TM^) manufacturer protocol for a 6-well plate. Similarly, SH-SY5Y and SKNF1 cells were stably transfected with a Cas9-expressing Piggybac plasmid (pPB[Exp]-TagBFP2/Neo-hPGK>hCas9) and a transposase plasmid (pRP[Exp]-mCherry-CAG>hyPBase) as described above. Stably transfected cells were then isolated by fluorescence-activated cell sorting (FACS).

### Cell Growth

SH-SY5Y and 3T3 cells expressing *GFP*, *H19*, or *DLK1* were seeded in 96-well plates with serum depleted medium (2.5% HI-FBS). The WST-8 cell proliferation assay kit (Cayman Chemical) was used to determine cell growth at 24-, 48-, and 72-hour time points. Growth rates were calculated relative to the 24-hour time point. For assays evaluating loss of *H19* or *DLK1*, SH-SY5Y cells transfected with siRNAs (as described below) were assayed for relative growth at 48 hours post-transfection. Cas9-expressing SH-SY5Y or SK-N-F1 were transfected with relevant gRNA constructs (as described below), and at 24 and 72 hours post-transfection, the level of fluorescence was measured. For all assays in the 96-well format, a seeding density of 5000 cells/well was used.

### Let-7 Target enrichment

SH-SY5Y and BE(2)C cells expressing *GFP* and H19 were plated in 6-well plates. After 48 hours, cells were harvested, and their derived cDNA was used in RT-qPCR (SYBR green system). Relative gene expression of *let-7* targets was determined using the 2^−ΔΔCt^ method. Primer sequences were available in Supplement Data File 11.

### Short interfering RNA (siRNA) Transfection

SH-SY5Y cells were seeded in 96-well plates and transfected with 1pmol siRNA according to Lipofectamine™ RNAiMAX transfection reagent (Invitrogen) manufacturer protocol at 48 hours post-transfection relative growth rate was measured. Also, CHLA-42 cells were seeded in 6 well plates and transfected with 25pmol H19 siRNAs according to Lipofectamine ™ RNAiMAX transfection reagent (Invitrogen) manufacturer protocol. Forty-eight hours post-transfection, cells were harvested, and RNA was isolated using the TRIzol^TM^ reagent. Subsequently, cDNA was synthesized using the Verso^TM^ cDNA kit. For all assays in 6 well format, a seeding density of 5x10^5^ cells/well. All of the siRNAs were acquired from IDT^TM^, Control siRNA sense: CGUUAAUCGCGUAUAAUACGCGUAT, antisense: AUACGCGUAUUAUACGCGAUUAACGAC; DLK1 siRNA 13.1 sense: CUCUUAAUGCAUGAUACAGAAUAAT, antisense AUUAUUCUGUAUCAUGCAUUAAGAGAG; DLK1 siRNA 13.2 sense: AAAAUGGAUUCUGCGAGGAUGACAA, antisense UUGUCAUCCUCGCAGAAUCCAUUUUGG; H19 siRNA 13.2 sense: AAAAUGGAUUCUGCGAGGAUGACAA, antisense UCCAGAGCCGAUUCCUGAGUCAGGUAG; H19 siRNA 13.3 sense : CAAGCAUUCCAUUACGCCCCAUCTC, antisense GAGAUGGGGCGUAAUGGAAUGCUUGAA.

### CRISPR/Cas-9 gene knockout

Cas9-expressing SH-SY5Y and SK-N-F1 *M-Non* NB cells were transfected with a single guide RNA (gRNA) constructs co-expressing either GFP and a single non-targeting gRNA or mCherry and single gRNAs targeting *RPL3*, *DLK1*, or *H19*. A third *H19*-targeting gRNA directly targets *miR-675* located within the first exon. Specifically, Cas9-expressing SH-SY5Y and SKNF1cells (100,000 cells/well) were transfected with 1 µg of appropriate gRNA vectors in 6-well plates on day 0. Cells were analyzed for percent fluorescence on days 1 and 3, and relative fluorescence on day 3 vs. day 1 was used to determine growth differences. Guide RNAs were designed as previously described^8^; sequences are available in Supplement Data File 11.

### Data Availability Statement

Accession codes for our NGS sequencing data (***Figure 4c***) will be provided prior to publication. Data for secondary analysis of the SEQC, Kocak, TARGET, and Oberthuer studies are publicly available on the r2 database (https://hgserver1.amc.nl/cgi-bin/r2/main.cgi). Data subsets used from these studies and experimental data we generated for growth assays (WST8 and gRNA/fluorescence) and qPCR analysis are provided in our Supporting Data Files along with our calculations and analyses.

## Acknowledgments

We would like to thank Dr. Diane K. Michelson for her expert assistance with the statistical analysis of genomic location-based differential gene expression. This study was supported by a Cancer Prevention and Research Institute of Texas Grant, RR180034 (PI J.T.P).

---

**Extended Data Figure 1.**
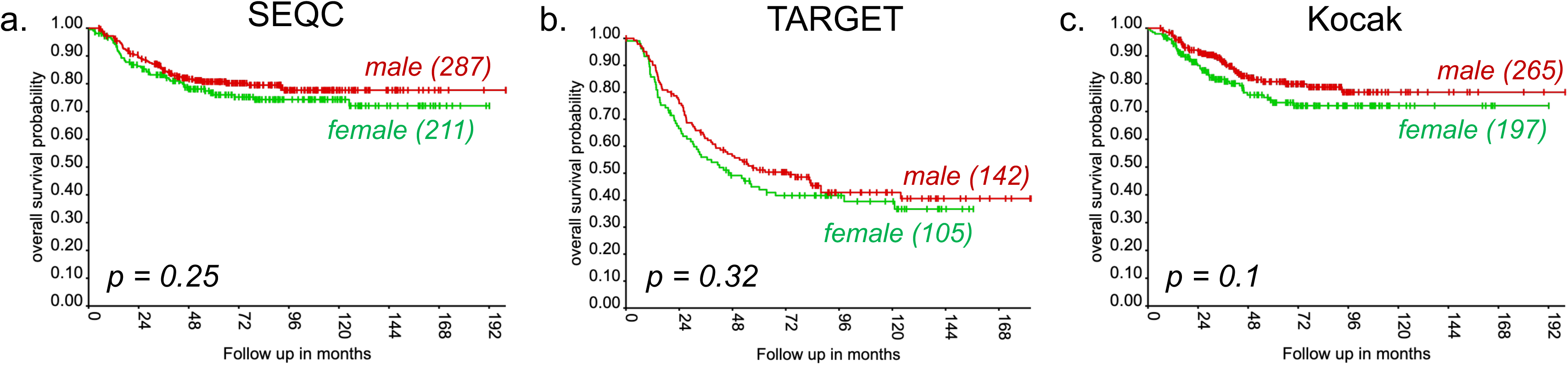
Overall survival probability of male and female neuroblastoma patients across datasets. OS in male and female NB from **a)** SEQC 498 (GSE62564) cohort, **b)** TARGET 249 (R2, Asgharzadeh) cohort, and **c)** Kocak 649 (GSE45547) cohort. P-values determined by log-rank test.

**Extended Data Figure 2.**
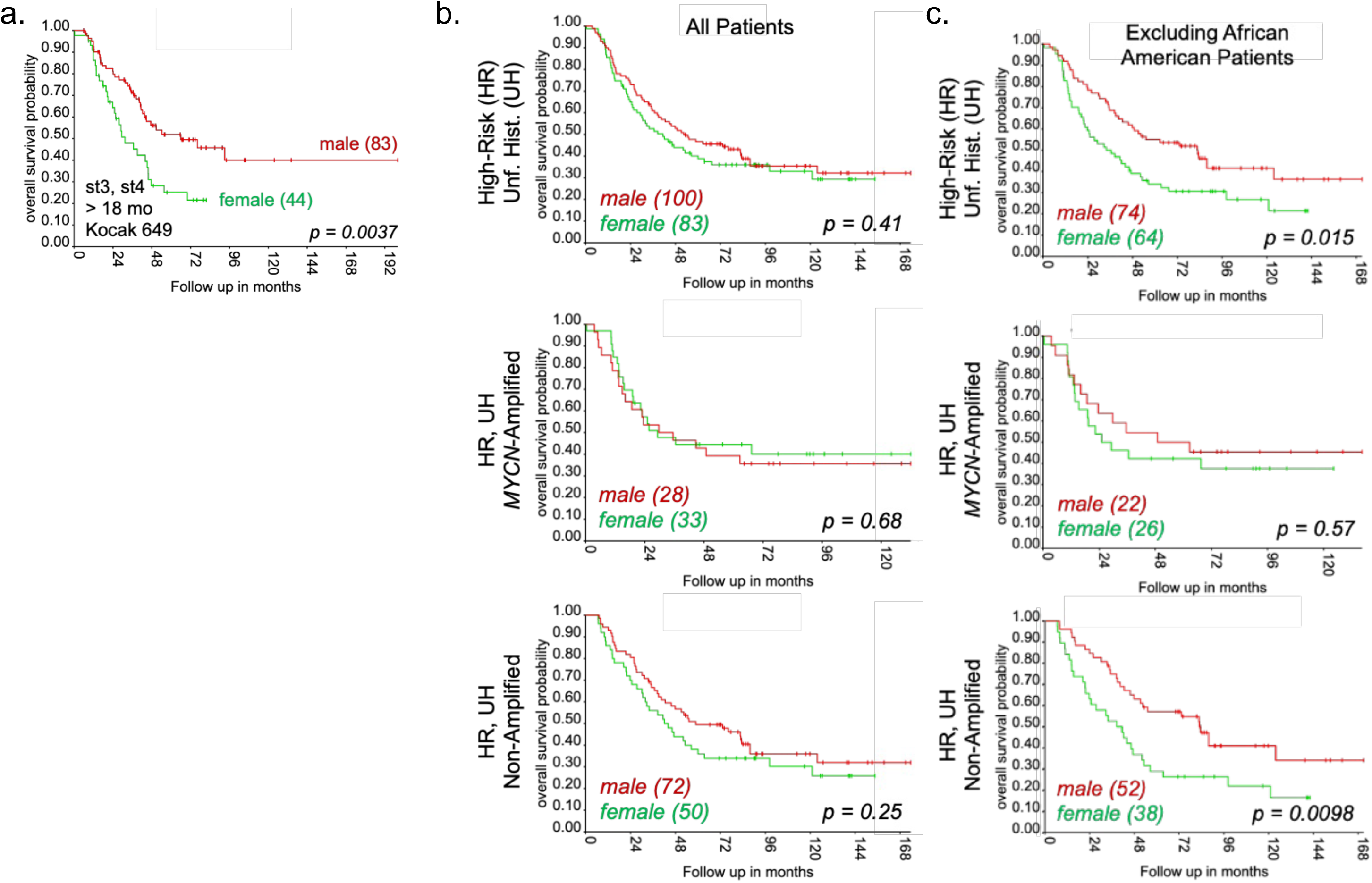
Overall survival of male and female neuroblastoma patients in additional patient studies. **a)** KM analysis of OS in male vs. female st 3 and 4 NB patients > 18 months old (approximating HR). Data source: Kocak 649 (GSE45547). **b)** KM analysis of OS in male and female *M-Amp*/HR, *M-Non*/HR neuroblastoma patients with unfavorable histology (UH). Subgroup determinants are detailed on y-axis for each graph; data source: TARGET 249 (R2, Asgharzadeh). **c)** KM analysis of OS in non-African American male and female HR, *M-Amp* and HR, *M-Non* neuroblastoma patients with unfavorable histology (UH). Subgroup determinants are detailed on y-axis for each graph; data source: TARGET 249 (R2, Asgharzadeh). All p-values were determined by the log-rank test.

**Extended Data Figure 3.**
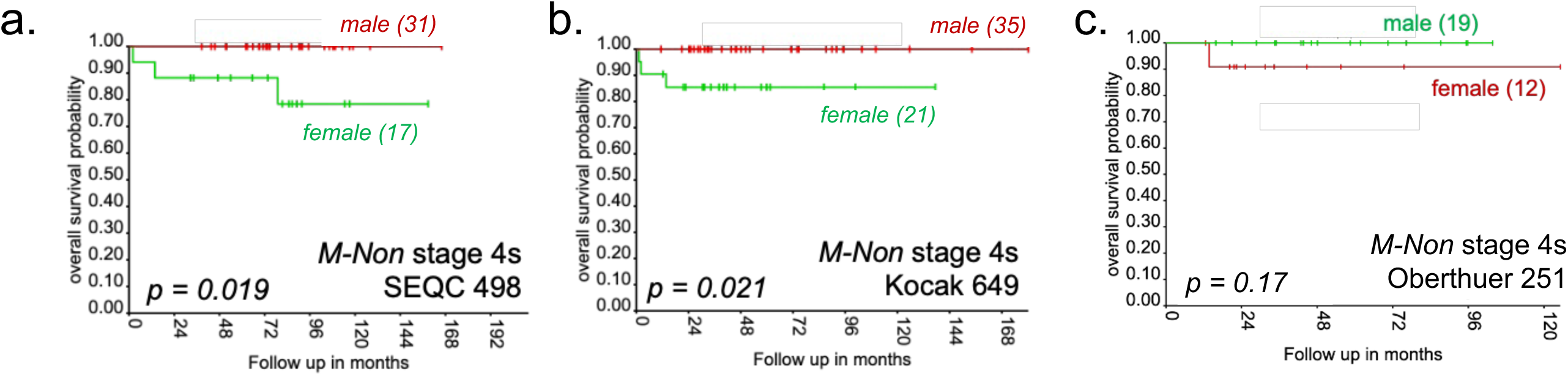
Sex-biased outcomes in M-Non, stage 4s neuroblastoma. OS in male and female NB patients from **a)** SEQC 498 (GSE62564) cohort, **b)** Kocak 649 (GSE45547) cohort, and **c)**. Oberthuer 251 (E-TABM-38) cohort. p values determined by log-rank test.

**Extended Data Figure 4.**
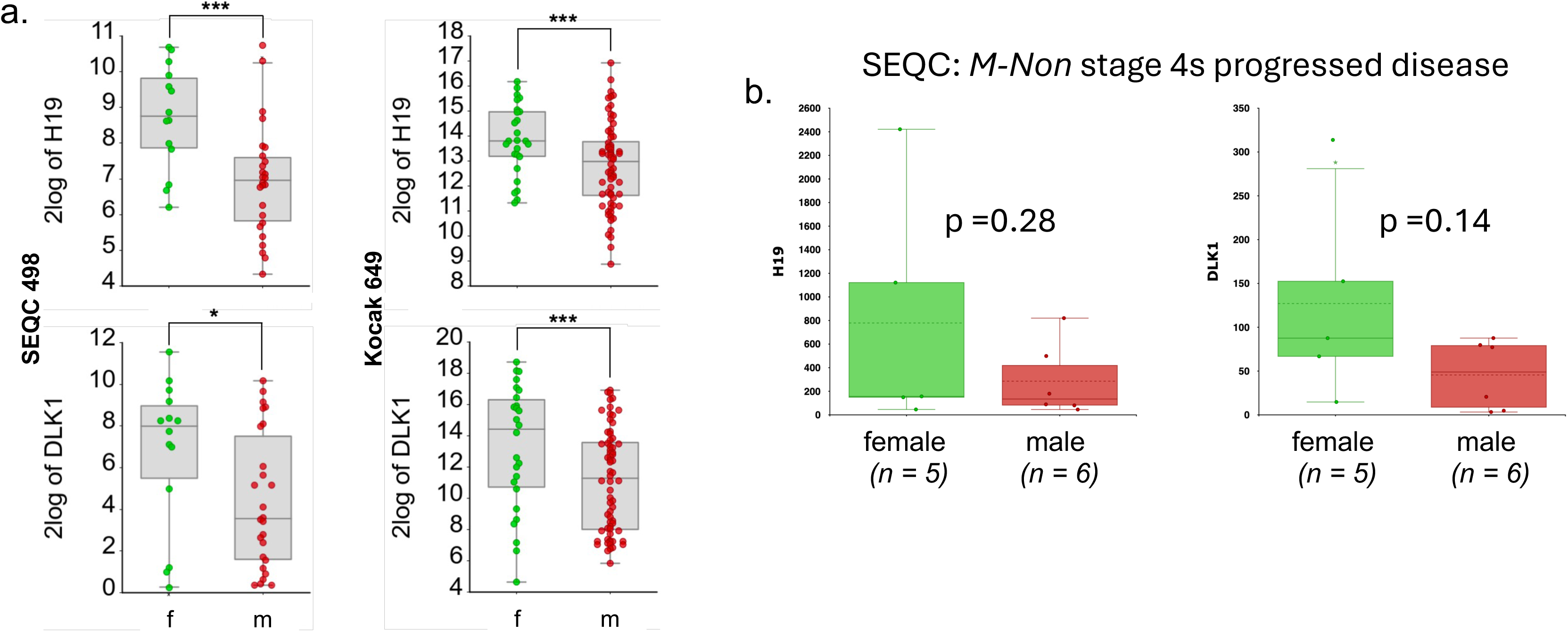
*H19* and *DLK1* expression in female and male M-Non/HR patients. **a)** Boxplots of *H19* (top) and *DLK1* (bottom) expression in male (m; red dots) vs. female (f; green dots) patients from subsets from two datasets: SEQC 498 (left; HR/*M-Non/*DFD subset; GSE62564) and Kocak 649 (right; *M-Non/*st4 and >18 months old subset; GSE45547). *p<0.05, ***p<0.001; determined by ANOVA. **b)** Boxplots of *H19* and *DLK1* expression in male and female M-Non stage4s progressed disease patients: Data source SEQC 498. p values determined by ANOVA

**Extended Data Figure 5.**
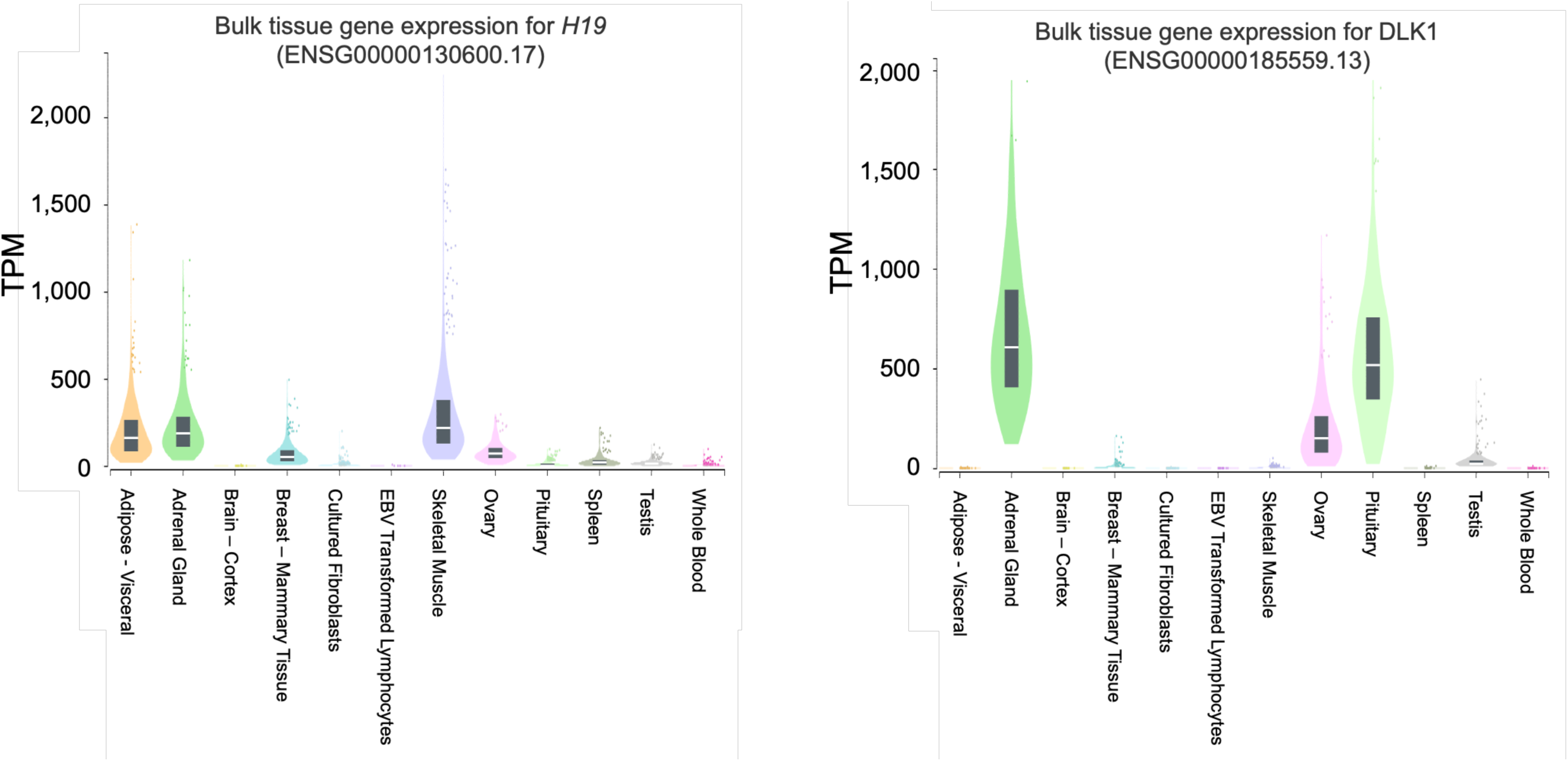
*H19* and *DLK1* expression levels in normal human tissues. Expression Values are shown in Transcripts per Million (TPM). Box plots are shown as median and 25th and 75th percentiles; outliers ± 1.5x the interquartile range. Data Source: GTEx Analysis Release V8 (dbGaP Accession phs000424.v8.p2)

**Extended Data Figure 6.**
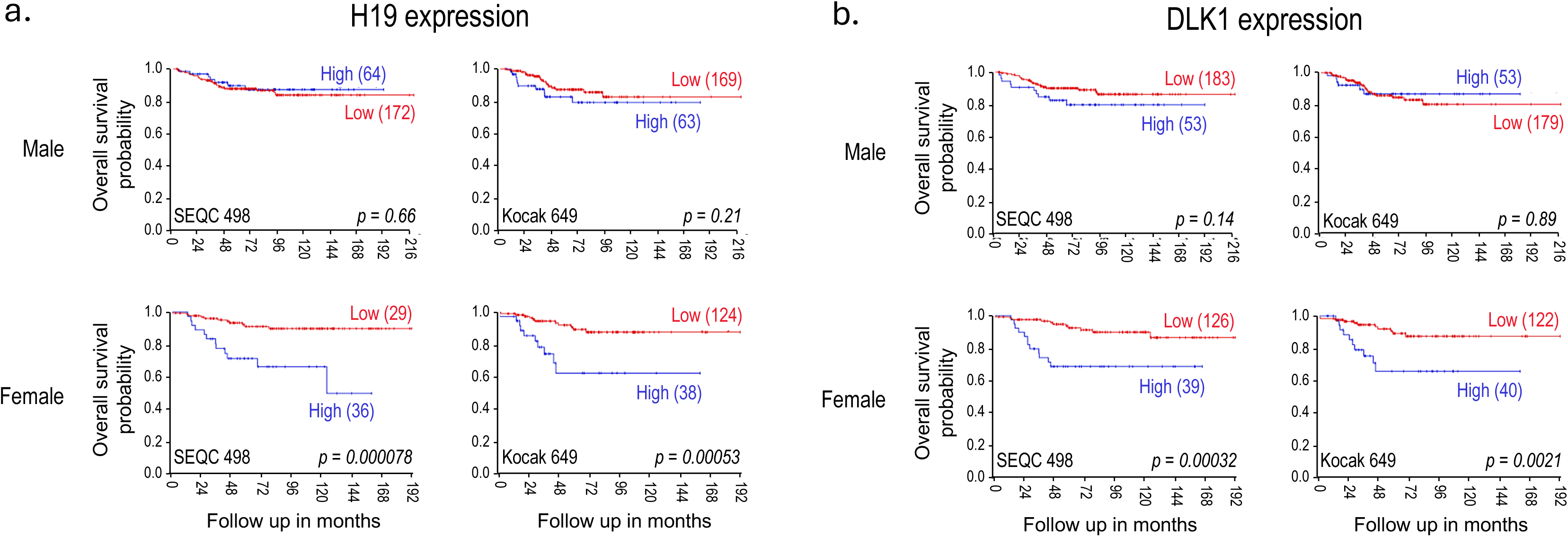
Overall survival in male and female *MYCN* non-amplified neuroblastoma patients relative to *H19* or *DLK1* expression. KM plots for males (top) and females (bottom) relative to expression (high or low) levels of **a)** *H19* or **b)** *DLK1* in *M-Non* NB. Data source: SEQC 498 (GSE62564) and Kocak 649 (GSE45547); source of the graph as listed on the respective figure. Red lines = low expression; blue lines = high expression; cutoff for high/low is average expression. p-values calculated by log-rank method.

**Extended Data Figure 7.**
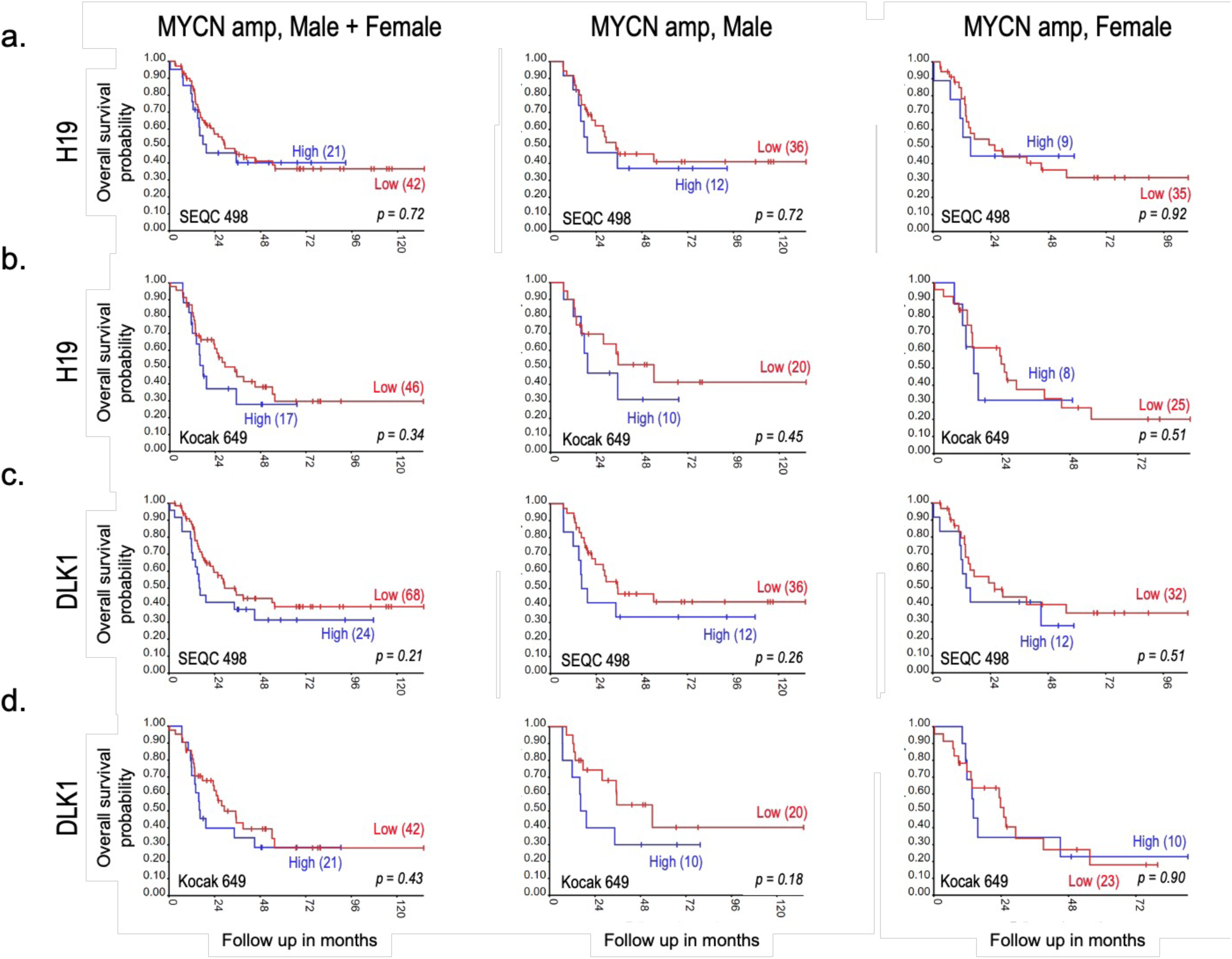
*H19* and *DLK1* are not associated with overall survival in *MYCN*-amplified disease. KM analaysis of OS in two cohorts (as labeled) of male and female *M-Amp* NB patients relative to **a,b)** *H19* or **c,d)** *DLK1* expression. In each panel, plots contain males + females (left), males only (center) and females only (right). Red line = low expression; blue line = high expression; high/low cutoff is average expression. Data source: SEQC 498 (GSE62564), Kocak 649 (GSE45547), R2; p-values calculated by log-rank method.

**Extended Data Figure 8.**
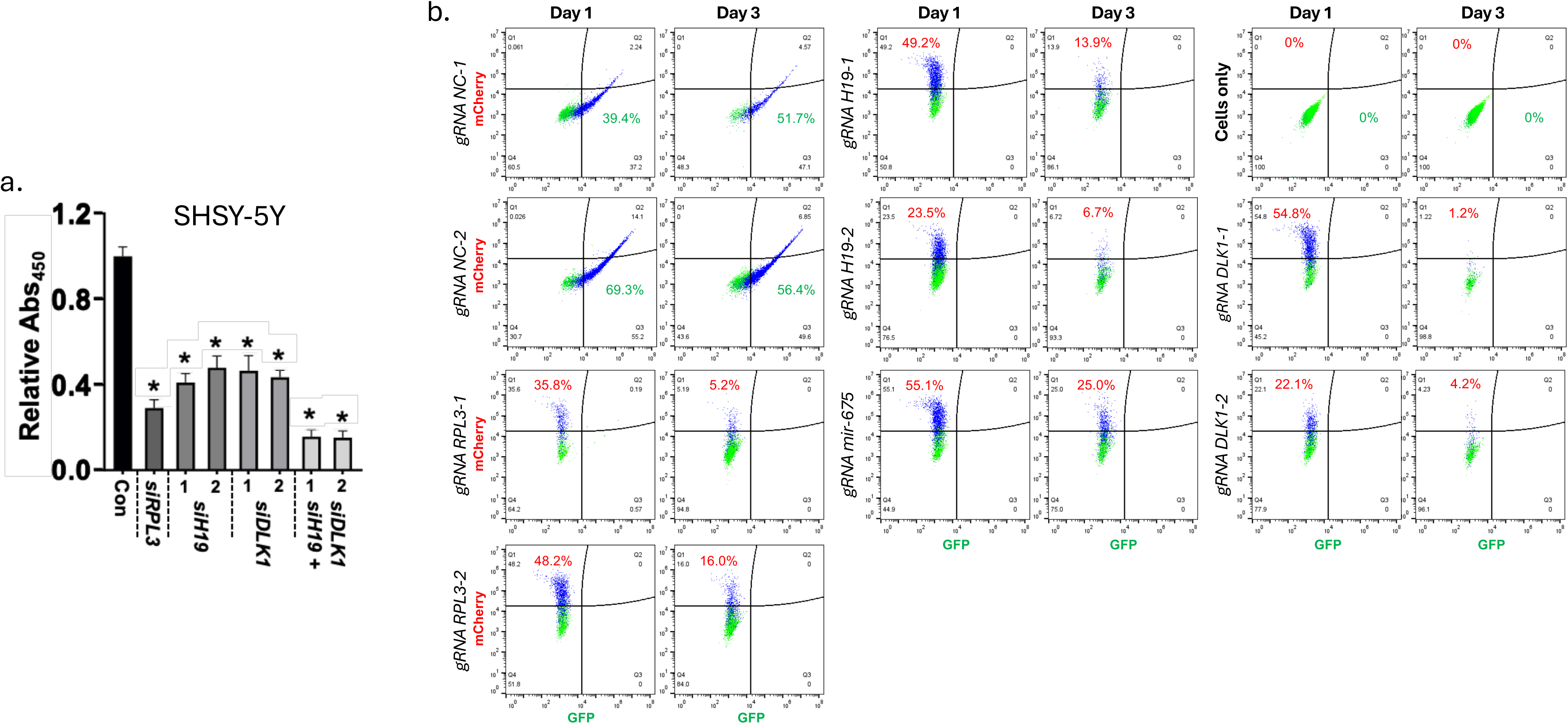
**a)** Growth effects of *H19* and *DLK1* knockdown in SHSY-5Y cells. Stated siRNAs were transfected on day 0 and assayed for growth (WST-8) 48 hours later. **b)** CRISPR/Cas9 mediated gene knockout of *H19* and *DLK1* in SHSY-5Y cells. Cells were transfected on d0 with control, *RPL3*, *DLK1* or *H19* targeting gRNAs. Loss of *H19* and *DLK1* targeted with gRNAs against negative control gRNA were measured at d1 and d3 via flow cytometry.

**Extended Data Figure 9.**
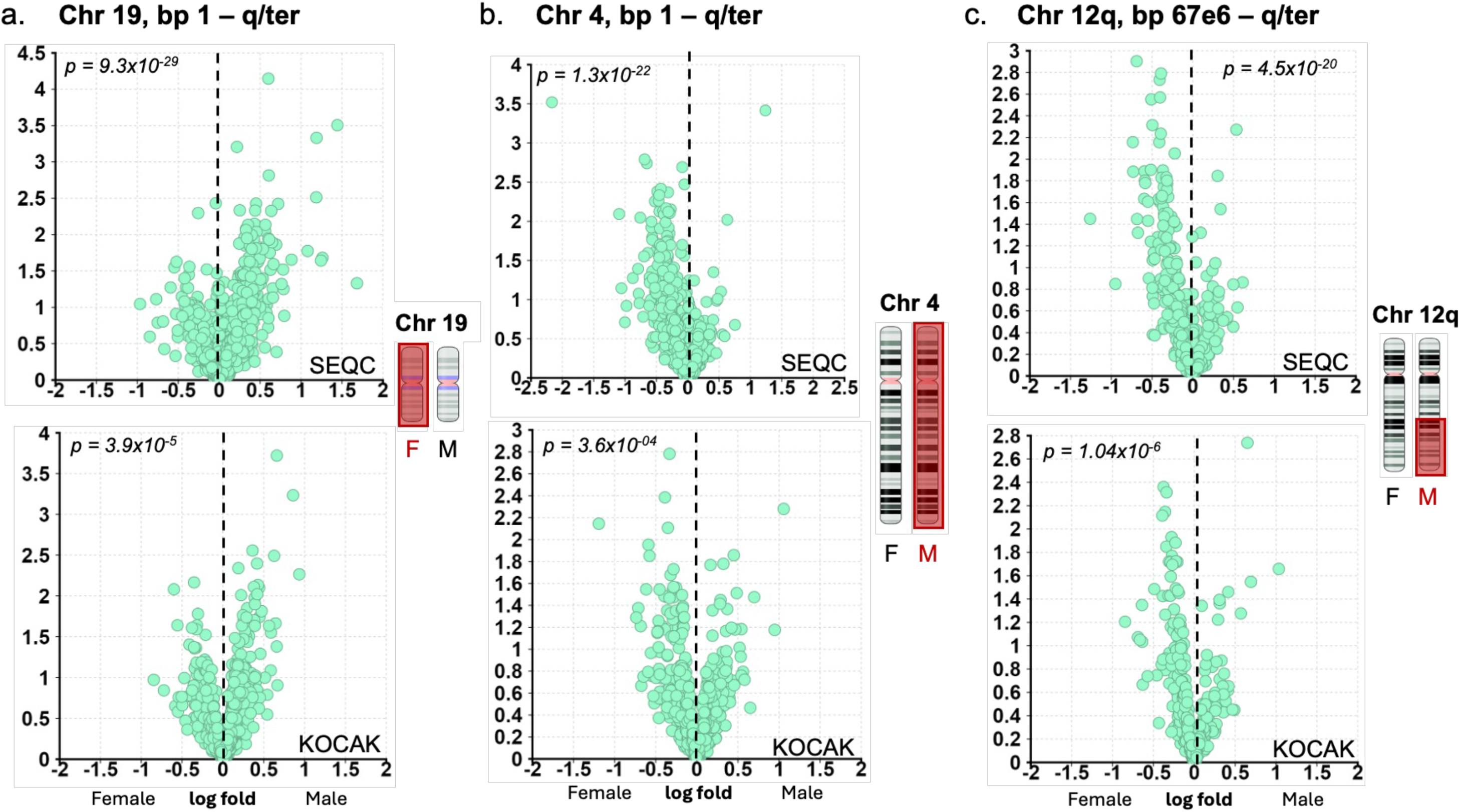
Additional chromosomal-linked gene expression data from male and female high-risk NB patients dead from disease. Volcano gene expression plots of male vs. female HR, NB patients from **a)** Chr 19, bp 1 – q/ter, and **b)** Chr 4, bp 1 – q/ter and **c)** Chr 12q, bp 67e6 – q/ter. In all panels, green dots = genes stemming from specified chromosome; red box over chromosome = relative chromosome deficiency in neuroblastoma. Data source: SEQC 498 (GSE62564, n=40) and Kocak 649 (GSE45547, n=94), as labeled.

**Extended Data Figure 10.**
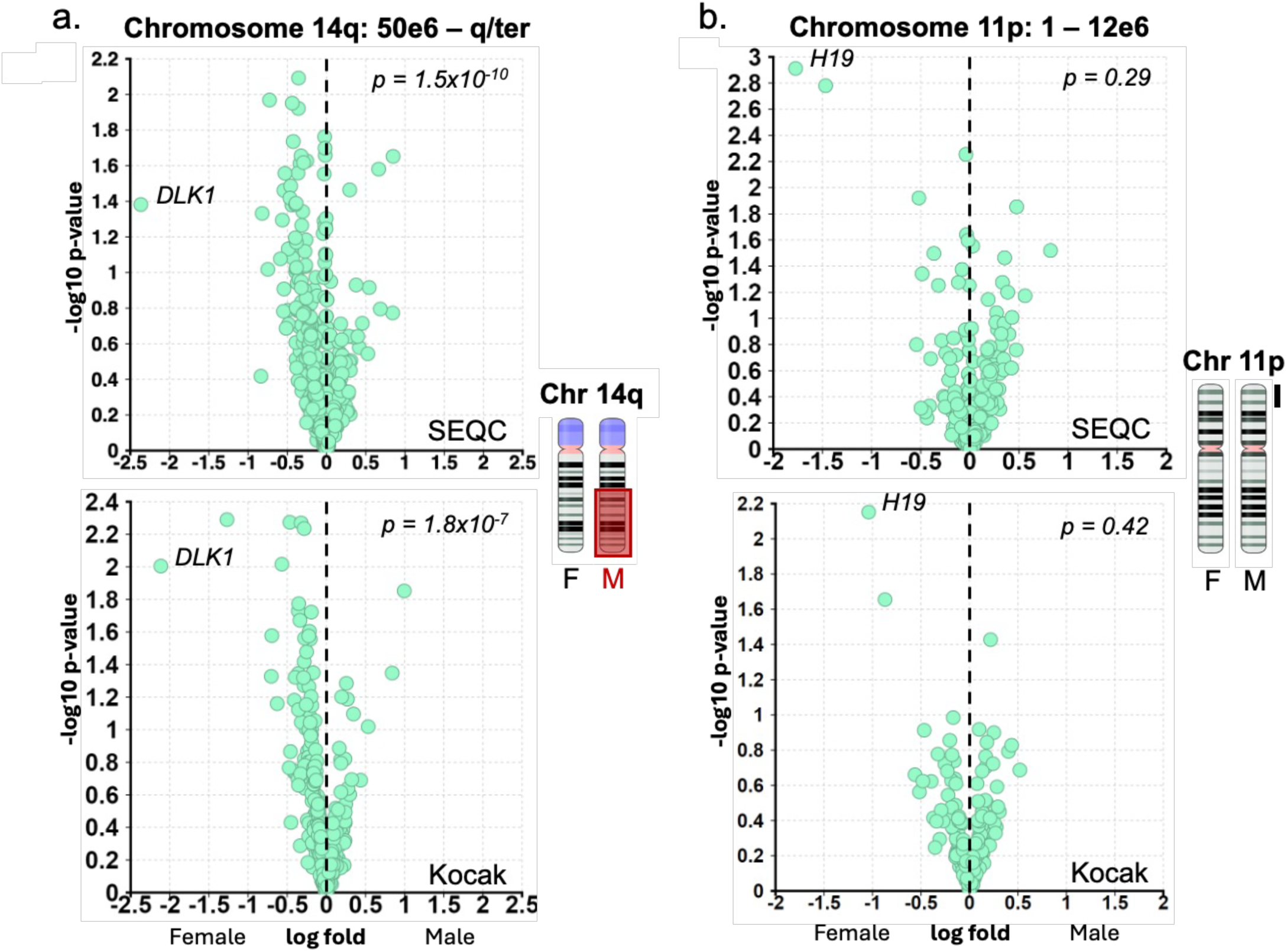
Volcano plots of genes located along **a)** Chr 14 50e6 -q/ter, and **b)** chr 11q 1 – 12e6 in male vs. female HR NB patients. In all panels, green dots = genes stemming from specified chromosome; red box over chromosome = relative chromosome deficiency in neuroblastoma. Data source as labeled on the graph: SEQC 498 (top; GSE62564, *M-Non*/HR/DFD subset; n=40) and Kocak 649 (Bottom; GSE45547, st4 patients >18 months old subset; n=94).

**Extended Data Figure 11.**
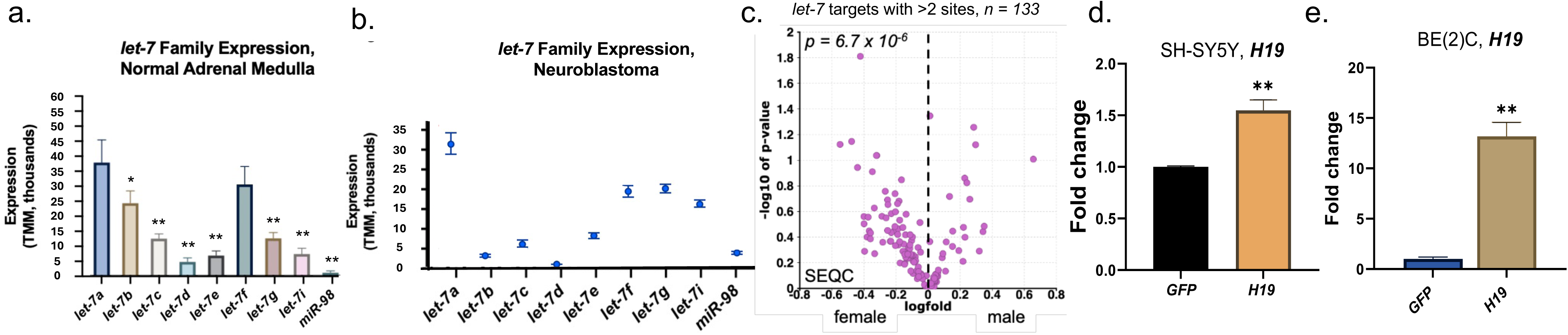
**a)** *Let-7* family member expression in the adrenal medulla, the tissue of origin for NB. Bars represent mean+/-standard deviation. Data source is GSE29742 including the following samples (n=6): GSM737418, GSM737419, GSM737420, GSM737421, GSM737422, GSM737423. p-values determined via Dunnett’s T3 Multiple Comparisons Test with *let-7a* as control; *p<0.05, **p<0.01; **b)** Mean expression of *let-7* family members in NB. Data source: Bell (GSE155945, n=97); error bars represent SEM. **c)** Volcano plot of differentially expressed *let-7* target mRNAs (containing >2 *let-7* sites) between female and male NB patients Data source: SEQC 498 (GSE62564; *M-Non*/HR/DFD subset; *n=40*). **d),e)** *H19* expression in SHSY-5Y, BE(2)C cells transfected with *H19* plasmid construct respectively (see *Methods*). **p<0.01

**Extended Data Figure 12.**
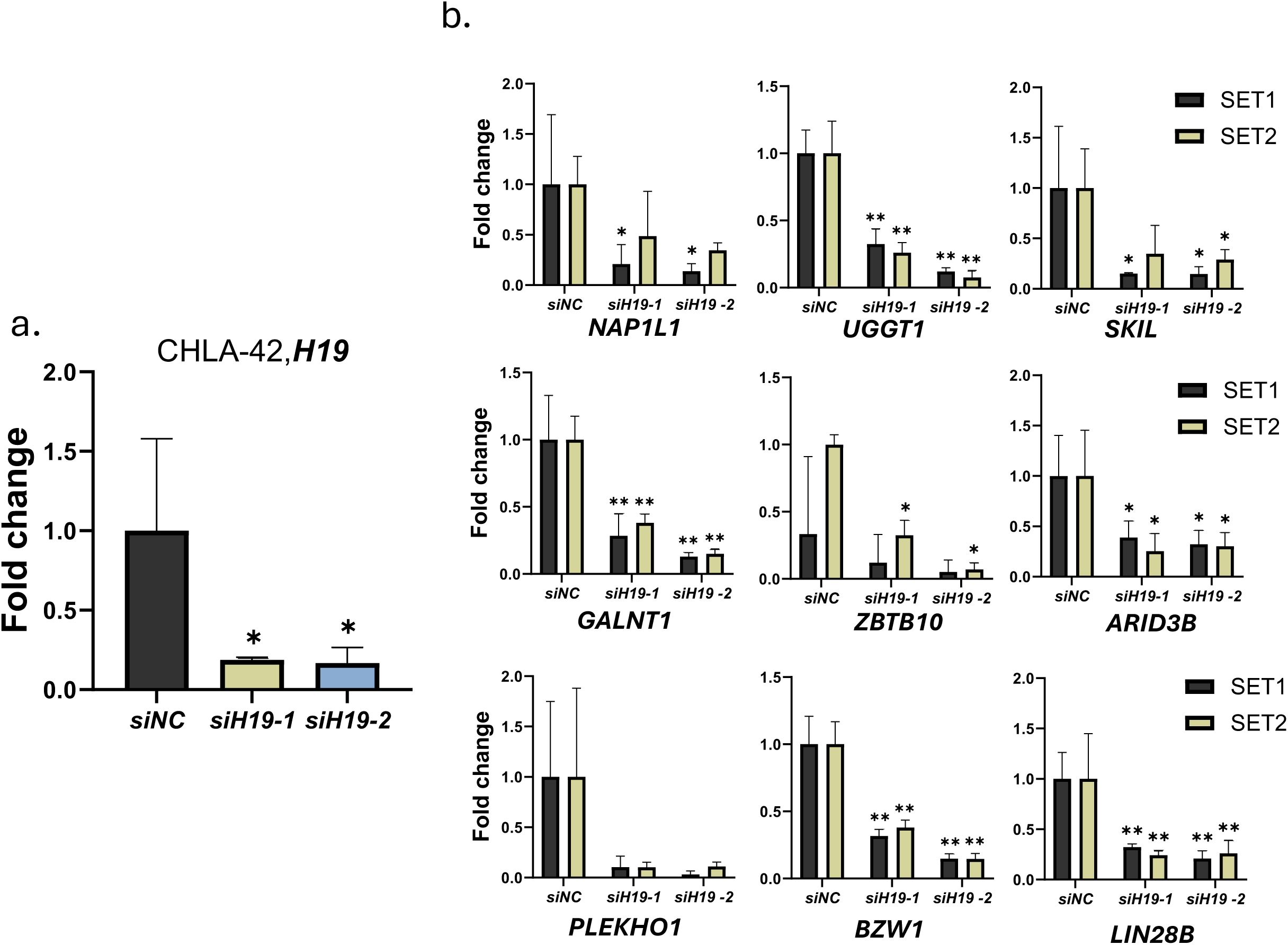
**a)** H19 expression in *M-Non* CHLA-42 cells in response to control siRNA or *H19* targeting siRNA transfection. **b)** Expression levels of *let-7* target genes in CHLA-42 cells, respectively, treated as in (a)Significance determined by two-way ANOVA, followed by Tukey’s multiple comparison tests; bars represent average +/- SEM; **p<0.01

**Extended Data Table 1.**
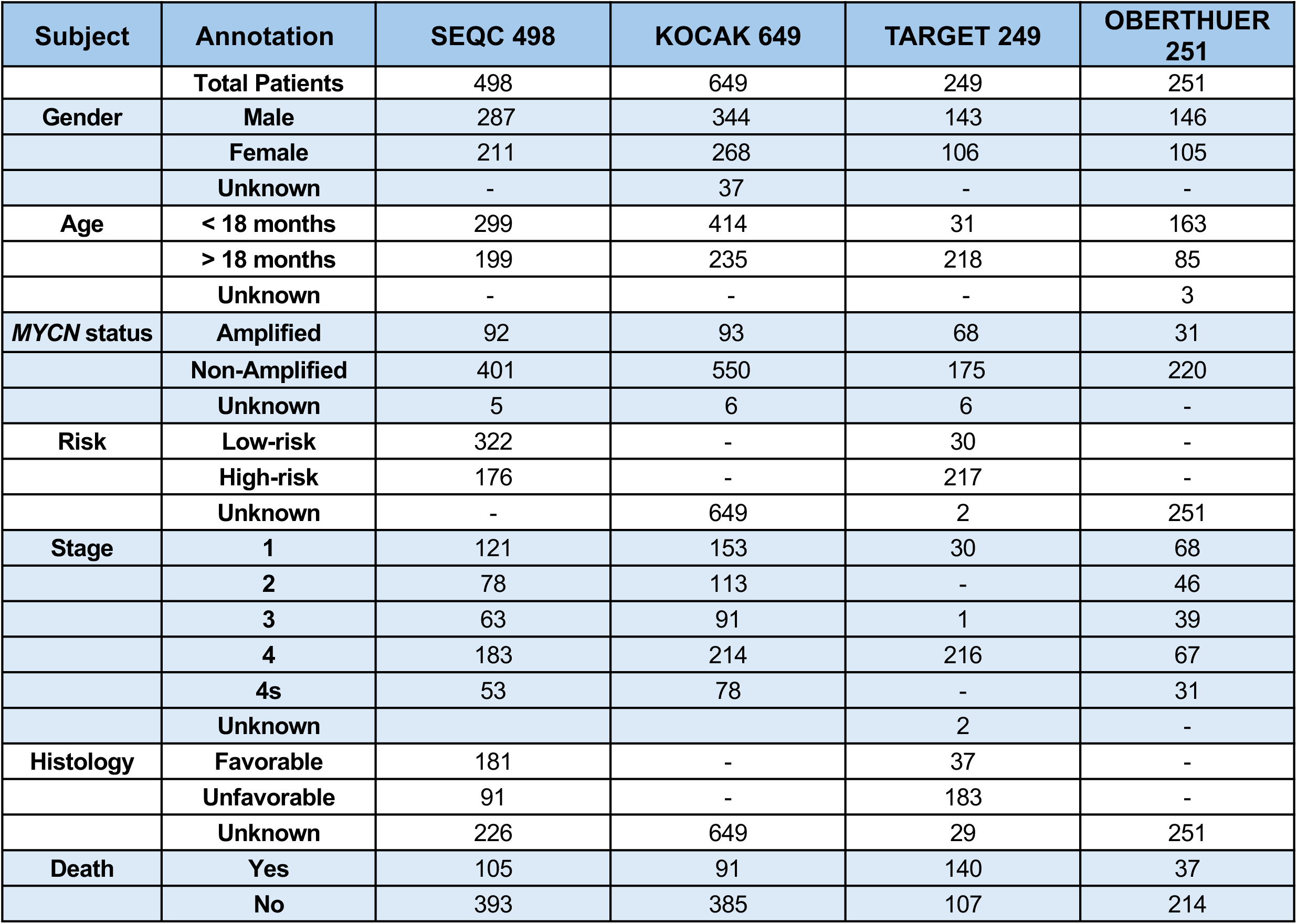
Neuroblastoma patient demographics and available annotations from the studies used in this manuscript.

## References

1. Irwin MS, Naranjo A, Zhang FF, et al. Revised Neuroblastoma Risk Classification System: A Report From the Children’s Oncology Group. Journal of Clinical Oncology. 2021;39(29):3229–3241. doi:10.1200/JCO.21.00278

2. DuBois SG, Macy, Margaret E., Henderson TO. High-Risk and Relapsed Neuroblastoma: Toward More Cures and Better Outcomes. American Society of Clinical Oncology Educational Book. 2022;(42):768–780. doi:10.1200/EDBK_349783

3. Williams LA, Spector LG. Survival Differences Between Males and Females Diagnosed With Childhood Cancer. JNCI Cancer Spectr. 2019;3(2). doi:10.1093/jncics/pkz032

4. Yan P, Qi F, Bian L, et al. Comparison of Incidence and Outcomes of Neuroblastoma in Children, Adolescents, and Adults in the United States: A Surveillance, Epidemiology, and End Results (SEER) Program Population Study. Med Sci Monit. 2020;26:e927218. doi:10.12659/MSM.927218

5. Maris JM, Guo C, White PS, et al. Allelic deletion at chromosome bands 11q14-23 is common in neuroblastoma. Med Pediatr Oncol. 2001;36(1):24–27. doi:10.1002/1096-911X(20010101)36:1<24::AID-MPO1007>3.0.CO;2-7

6. Attiyeh E, London W, Mosse YP, et al. Chromosome 1p and 11q deletions and outcome in neuroblastoma. New England Journal of Medicine. 2005;353(21):2243–2253. Accessed November 13, 2014. http://www.nejm.org/doi/full/10.1056/nejmoa052399

7. Kallen AN, Zhou XB, Xu J, et al. The Imprinted H19 LncRNA Antagonizes Let-7 MicroRNAs. Mol Cell. 2013;52(1):101–112. doi:10.1016/j.molcel.2013.08.027

8. Powers JT, Tsanov KM, Pearson DS, et al. Multiple mechanisms disrupt the let-7 microRNA family in neuroblastoma. Nature. 2016;535(7611):246-251. doi:10.1038/nature18632

9. Johnsen JI, Dyberg C, Wickström M. Neuroblastoma—A Neural Crest Derived Embryonal Malignancy. Front Mol Neurosci. 2019;12. doi:10.3389/fnmol.2019.00009

10. Tsubota S, Kadomatsu K. Origin and mechanism of neuroblastoma. Oncoscience. 2017;4(7-8):70–72. doi:10.18632/oncoscience.360

11. Liang WH, Federico SM, London WB, et al. Tailoring Therapy for Children With Neuroblastoma on the Basis of Risk Group Classification: Past, Present, and Future. JCO Clin Cancer Inform. 2020;(4):895–905. doi:10.1200/CCI.20.00074

12. Irwin MS, Naranjo A, Zhang FF, et al. Revised Neuroblastoma Risk Classification System: A Report From the Children’s Oncology Group. Journal of Clinical Oncology. 2021;39(29):3229–3241. doi:10.1200/JCO.21.00278

13. Tas ML, Nagtegaal M, Kraal KCJM, et al. Neuroblastoma stage 4S: Tumor regression rate and risk factors of progressive disease. Pediatr Blood Cancer. 2020;67(4). doi:10.1002/pbc.28061

14. Nickerson HJ, Matthay KK, Seeger RC, et al. Favorable Biology and Outcome of Stage IV-S Neuroblastoma With Supportive Care or Minimal Therapy: A Children’s Cancer Group Study. Journal of Clinical Oncology. 2000;18(3):477–477. doi:10.1200/JCO.2000.18.3.477

15. Colon NC, Chung DH. Neuroblastoma. Adv Pediatr. 2011;58(1):297–311. doi:10.1016/j.yapd.2011.03.011

16. Cohn SL, Pearson ADJ, London WB, et al. The International Neuroblastoma Risk Group (INRG) classification system: an INRG Task Force report. J Clin Oncol. 2009;27(2):289–297. doi:10.1200/JCO.2008.16.6785

17. Otte J, Dyberg C, Pepich A, Johnsen JI. MYCN Function in Neuroblastoma Development. Front Oncol. 2021;10. doi:10.3389/fonc.2020.624079

18. He Y, Su Y, Zeng J, et al. Cancer-specific survival after diagnosis in men versus women: A pan-cancer analysis. MedComm (Beijing). 2022;3(3):e145. doi:10.1002/mco2.145

19. Lopes-Ramos CM, Quackenbush J, DeMeo DL. Genome-Wide Sex and Gender Differences in Cancer. Front Oncol. 2020;10. doi:10.3389/fonc.2020.597788

20. Zavala VA, Bracci PM, Carethers JM, et al. Cancer health disparities in racial/ethnic minorities in the United States. Br J Cancer. 2021;124(2):315–332. doi:10.1038/s41416-020-01038-6

21. Holowatyj AN, Wen W, Gibbs T, et al. Racial/Ethnic and Sex Differences in Somatic Cancer Gene Mutations among Patients with Early-Onset Colorectal Cancer. Cancer Discov. 2023;13(3):570–579. doi:10.1158/2159-8290.CD-22-0764

22. Breen CJ, O’Meara a, McDermott M, Mullarkey M, Stallings RL. Coordinate Deletion of Chromosome 3p and 11q in Neuroblastoma Detected by Comparative Genomic Hybridization. Cancer Genet Cytogenet. 2000;120(1):44–49. doi:10.1016/S0165-4608(99)00252-6

23. Theissen J, Oberthuer A, Hombach A, et al. Chromosome 17 / 17q Gain and Unaltered Profiles in High Resolution Array-CGH are Prognostically Informative in Neuroblastoma. 2014;649(April):639–649. doi:10.1002/gcc

24. Roush S, Slack FJ. The let-7 family of microRNAs. Trends Cell Biol. 2008;18(10). doi:10.1016/j.tcb.2008.07.007

25. Perini G, Milazzo G, Narayan N, Ekert PG. Letting the breaks off MYCN. Cell Death Differ. Published online 2016:1–2. doi:10.1038/cdd.2016.112

26. Pittaway JFH, Lipsos C, Mariniello K, Guasti L. The role of delta-like non-canonical Notch ligand 1 (DLK1) in cancer. Endocr Relat Cancer. 2021;28(12):R271–R287. doi:10.1530/ERC-21-0208

27. Grassi ES, Pietras A. Emerging Roles of DLK1 in the Stem Cell Niche and Cancer Stemness. Journal of Histochemistry & Cytochemistry. 2022;70(1):17–28. doi:10.1369/00221554211048951

28. Yang J, Qi M, Fei X, Wang X, Wang K. LncRNA H19: A novel oncogene in multiple cancers. Int J Biol Sci. 2021;17(12):3188–3208. doi:10.7150/ijbs.62573

29. Wu B, Zhang Y, Yu Y, et al. Long Noncoding RNA H19: A Novel Therapeutic Target Emerging in Oncology Via Regulating Oncogenic Signaling Pathways. Front Cell Dev Biol. 2021;9. doi:10.3389/fcell.2021.796740

30. Campbell K, Siegel DA, Umaretiya PJ, et al. A comprehensive analysis of neuroblastoma incidence, survival, and racial and ethnic disparities from 2001 to 2019. Pediatr Blood Cancer. 2024;71(1). doi:10.1002/pbc.30732

31. SEQC/MAQC-III Consortium. A comprehensive assessment of RNA-seq accuracy, reproducibility and information content by the Sequencing Quality Control Consortium. Nat Biotechnol. 2014;32(9):903–914. doi:10.1038/nbt.2957

32. Kocak H, Ackermann S, Hero B, et al. Hox-C9 activates the intrinsic pathway of apoptosis and is associated with spontaneous regression in neuroblastoma. Cell Death Dis. 2013;4(4):e586. doi:10.1038/cddis.2013.84

33. Grossman RL, Heath AP, Ferretti V, et al. Toward a Shared Vision for Cancer Genomic Data. New England Journal of Medicine. 2016;375(12):1109–1112. doi:10.1056/NEJMp1607591

34. Kawano A, Hazard FK, Chiu B, et al. Stage 4S Neuroblastoma. American Journal of Surgical Pathology. 2021;45(8):1075–1081. doi:10.1097/PAS.0000000000001647

35. White PS, Thompson PM, Gotoh T, et al. Definition and characterization of a region of 1p36.3 consistently deleted in neuroblastoma. Oncogene. 2005;24(16):2684–2694. doi:10.1038/sj.onc.1208306

36. Juan Ribelles A, Gargallo P, Ferriol C, et al. Distribution of segmental chromosomal alterations in neuroblastoma. Clinical and Translational Oncology. 2021;23(6):1096–1104. doi:10.1007/s12094-020-02497-2

37. Janoueix-Lerosey I, Schleiermacher G, Michels E, et al. Overall Genomic Pattern Is a Predictor of Outcome in Neuroblastoma. Journal of Clinical Oncology. 2009;27(7):1026–1033. doi:10.1200/JCO.2008.16.0630

38. Mlakar V, Jurkovic Mlakar S, Lopez G, Maris JM, Ansari M, Gumy-Pause F. 11q deletion in neuroblastoma: a review of biological and clinical implications. Mol Cancer. 2017;16(1):114. doi:10.1186/s12943-017-0686-8

39. Caron H, van Sluis P, Buschman R, et al. Allelic loss of the short arm of chromosome 4 in neuroblastoma suggests a novel tumour suppressor gene locus. Hum Genet. 1996;97(6):834–837. doi:10.1007/BF02346199

40. Su WT, Alaminos M, Mora J, Cheung NK, La Quaglia MP, Gerald WL. Positional gene expression analysis identifies 12q overexpression and amplification in a subset of neuroblastomas. Cancer Genet Cytogenet. 2004;154(2):131–137. doi:10.1016/j.cancergencyto.2004.02.009

41. Martinez-Monleon A, Kryh Öberg H, Gaarder J, et al. Amplification of CDK4 and MDM2: a detailed study of a high-risk neuroblastoma subgroup. Sci Rep. 2022;12(1):12420. doi:10.1038/s41598-022-16455-1

42. Roy N Van, Forus A, Myklebost O, Cheng NC, Versteeg R, Speleman F. Identification of two distinct chromosome 12-derived amplification units in neuroblastoma cell line NGP. Cancer Genet Cytogenet. 1995;82(2):151–154. doi:10.1016/0165-4608(95)00034-M

43. Peng F, Li TT, Wang KL, et al. H19/let-7/LIN28 reciprocal negative regulatory circuit promotes breast cancer stem cell maintenance. Cell Death Dis. 2017;8(1):e2569–e2569. doi:10.1038/cddis.2016.438

44. Peng F, Li TT, Wang KL, et al. H19/let-7/LIN28 reciprocal negative regulatory circuit promotes breast cancer stem cell maintenance. Cell Death Dis. 2017;8(1):e2569–e2569. doi:10.1038/cddis.2016.438

45. Peng F, Wang JH, Fan WJ, et al. Glycolysis gatekeeper PDK1 reprograms breast cancer stem cells under hypoxia. Oncogene. 2018;37(8):1062–1074. doi:10.1038/onc.2017.368

46. Su H, Xu X, Yan C, et al. LncRNA H19 promotes the proliferation of pulmonary artery smooth muscle cells through AT1R via sponging let-7b in monocrotaline-induced pulmonary arterial hypertension. Respir Res. 2018;19(1):254. doi:10.1186/s12931-018-0956-z

47. Lopes-Ramos CM, Quackenbush J, DeMeo DL. Genome-Wide Sex and Gender Differences in Cancer. Front Oncol. 2020;10. doi:10.3389/fonc.2020.597788

48. He Y, Su Y, Zeng J, et al. Cancer-specific survival after diagnosis in men versus women: A pan-cancer analysis. MedComm (Beijing). 2022;3(3). doi:10.1002/mco2.145

49 R2: Genomics Analysis and Visualization Platform (http://r2.amc.nl).

50. Lecerf C, Le Bourhis X, Adriaenssens E. The long non-coding RNA H19: an active player with multiple facets to sustain the hallmarks of cancer. Cellular and Molecular Life Sciences. 2019;76(23):4673–4687. doi:10.1007/s00018-019-03240-z

51. Yu H, Li S, Wu S xiong, Huang S, Li S, Ye L. The prognostic value of long non-coding RNA H19 in various cancers. Medicine. 2020;99(2):e18533. doi:10.1097/MD.0000000000018533

52. Astuti D, Latif F, Wagner K, et al. Epigenetic alteration at the DLK1-GTL2 imprinted domain in human neoplasia: analysis of neuroblastoma, phaeochromocytoma and Wilms’ tumour. Br J Cancer. 2005;92(8):1574–1580. doi:10.1038/sj.bjc.6602478

53. Fukuzawa R, Heathcott RW, Morison IM, Reeve AE. Imprinting, expression, and localisation of DLK1 in Wilms tumours. J Clin Pathol. 2005;58(2):145–150. doi:10.1136/jcp.2004.021717

54. Pittaway JFH, Lipsos C, Mariniello K, Guasti L. The role of delta-like non-canonical Notch ligand 1 (DLK1) in cancer. Endocr Relat Cancer. 2021;28(12):R271–R287. doi:10.1530/ERC-21-0208

55. Alipoor B, Parvar SN, Sabati Z, Ghaedi H, Ghasemi H. An updated review of the H19 lncRNA in human cancer: molecular mechanism and diagnostic and therapeutic importance. Mol Biol Rep. 2020;47(8):6357–6374. doi:10.1007/s11033-020-05695-x

56. Sellers ZP, Schneider G, Maj M, Ratajczak MZ. Analysis of the Paternally-Imprinted DLK1–MEG3 and IGF2– H19 Tandem Gene Loci in NT2 Embryonal Carcinoma Cells Identifies DLK1 as a Potential Therapeutic Target. Stem Cell Rev Rep. 2018;14(6):823–836. doi:10.1007/s12015-018-9838-5

57. Takada S. Epigenetic analysis of the Dlk1-Gtl2 imprinted domain on mouse chromosome 12: implications for imprinting control from comparison with Igf2-H19. Hum Mol Genet. 2002;11(1):77–86. doi:10.1093/hmg/11.1.77

